# Density-dependent feedback and higher-order interactions enable coexistence in phage-bacteria community dynamics

**DOI:** 10.1101/2025.05.15.651590

**Authors:** Raunak Dey, Ashley R. Coenen, Natalie E. Solonenko, Marie N. Burris, Anna I. Mackey, Julia Galasso, Christine L. Sun, David Demory, Daniel Muratore, Stephen J. Beckett, Matthew B. Sullivan, Joshua S. Weitz

## Abstract

Diverse phage-bacteria communities coexist at high densities in environmental, agricultural, and human-associated microbiomes. Phage-bacteria coexistence is often attributed to coevolutionary processes mediated by complex, pairwise infection networks. Here, using *in vitro* experiments and mathematical models, we explore how higher-order interactions function as a complementary, ecological feedback mechanism to stabilize phage-bacteria communities. To do so, we examine an environmentally-derived, synthetic phage-bacteria community comprised of five marine heterotrophic bacteria (*Cellulophaga baltica* and *Pseudoalteromonas* strains) and five associated phage. We used Bayesian inference to reconstruct free phage production in one-step growth experiments and then forecasted pairwise phage-bacteria community dynamics over multiple infection cycles. In contrast to model predictions of rapid bacterial population collapse, each bacterial strain persisted in the community. We hypothesized and then experimentally validated the relevance of infection attenuation at relatively high viral densities. We extended models into a community context, corroborating complex coexistence of all phage and bacteria. Life history traits inferred in community fits often differed from those inferred in a pairwise context, implicating higher-order interactions as an additional, ecological stabilization mechanism. Follow-up experiments confirm that phage traits (including burst size) can shift when infecting single vs. multiple strains. More broadly, these findings suggest that complex community coexistence of phage and bacteria may be more common than anticipated when including feedback mechanisms outside of the growth-dominated regimes of fitted pairwise models that do not reflect the full scope of ecologically relevant contexts.

## 1 Main

Bacteriophage (‘phage’) shape bacterial population dynamics in human-associated, built environments, and environmental microbiomes^1–11^. Phage infections of bacteria are highly specific, such that realized interactions between phage and bacteria can be represented as complex, sparse networks^12^. At microevolutionary scales, phage-bacteria coevolve and often generate interaction networks with nested structures such that phage span a range of generalists to specialists and bacteria span a range of permissive to resistant^7^. At larger scales, phage-bacteria interaction networks can be characterized in terms of modules in which a subset of phage preferentially infect restricted subsets of bacteria^8^. Phage-bacteria interaction networks can also exhibit multi-scale structure, e.g., both in large-scale marine surveys ^8,10^ and within experimental coevolutionary studies beginning with joint inoculation of a single phage and bacterial type^13^. However, it remains an open challenge to connect pairwise interaction network structure with ecological feedback mechanisms that enable the emergence and maintenance of complex phage-bacteria communities.

Characterizing pairwise interactions is a first step in developing predictive models of community dynamics. For virus-microbe interactions, doing so requires measurements of associated life history traits. Baseline interactions between phage and bacteria can be characterized in a pairwise fashion given cultured isolates^7,13,14^. Ecologically relevant life history traits include bacterial growth rates and viral adsorption rates, latent periods, and burst sizes^15–17^. Scaling up pairwise interactions to community interactions has been used to predict coarse-grained patterns and processes of microbial community assemblages^18–21^. However, pairwise interactions often fail to capture the emergent stability and co-existence of communities that arise from factors such as higher-order interactions^22–25^, context-dependent emergent effects^26–28^, temporal partitioning^29^, density-dependent feedback^30^, and environmental feedback^31^. Pairwise models typically assume static interactions, ignoring the effects of changing environmental conditions and feedback loops^32–34^. These mechanisms may also be relevant in reconciling gaps between predicted community outcomes derived from pairwise interactions between phage and bacteria.

In this paper, we combine ecological theory, models, and direct experimental measurements to assess the limits to predictability of dynamics within a community of intermediate complexity given five bacterial strains from two different species of marine heterotrophic bacteria (*Cellulophaga baltica* and *Pseudoalteromonas* strains) and five virulent bacteriophage^35–37^. All five bacterial strains and five phage strains were originally isolated from similar conditions in marine waters and coexist in a synthetic community in laboratory conditions. Beginning with pairwise interactions, we used a nonlinear population model to infer life history traits from short, time series measurements of phage and bacteria abundance. The nonlinear model predicts rapid bacterial collapse within a few lytic cycles – in direct contrast to experimental evidence from longer time series. Through a process of model-data integration, we find that higher-order interactions and density-dependent feedback can enable phage-bacteria coexistence in complex communities. This phage-bacteria coexistence was not anticipated from models parameterized based on short-term, pairwise experiments in regimes where viral densities are relatively low. As we show, this iterative use of models and experiments reveals mechanisms that may enable coexistence of complex phage and bacterial communities – and provides a caution for extrapolating community-scale outcomes from a narrower scope of conditions than expected in ecologically relevant contexts.

## 2 Methods

### Strains and growth conditions

Five bacterial strains were used: *Cellulophaga baltica* strains NN016038, #4 (1), and #18 (2) were isolated from the Baltic Sea in 1994 (NN016038, hereafter CBA 38) and (3) in 2000 (#4 and #18, hereafter CBA 4 and CBA 18^38^); and *Pseudoalteromonas* sp.H100 (4) and 13-15 (5) were isolated from the North Sea in 1990 (hereafter PSA H100 and PSA 13-15^39^). Five bacteriophages were used: 1) *ϕ* 38:1, a podovirus isolated from the Baltic Sea in 2005 on CBA 38, 2) *ϕ* 18:2, a siphovirus isolated from the Baltic Sea in 2000 on CBA 18, 3) *ϕ* 18:3, a podovirus isolated from the Baltic Sea in 2005 on CBA 18^38^, 4) PSA-HP1, a podovirus isolated from the North Sea in 1990 on PSA H100, and 5) PSA-HS6, a siphovirus isolated from the North Sea in 1990 on PSA 11-68 (strain not included in this study^40^). Bacteria were grown on *Pseudoalteromonas-Cellulophaga* (PC) plates (20.5g Sigma Sea Salts, 1.5g peptone, 1.5g proteose peptone, 0.6g yeast extract, 13g agar/L) at room temperature (RT). Single colonies were inoculated and grown stationary at RT overnight in PC liquid growth medium (20.5g Sigma Sea Salts, 0.75g peptone, 0.5g yeast extract, 0.25g proteose peptone, 0.25g casamino acids, 1.5ml glycerol/L). Phage strains were stored in phage buffer (20.5g Sigma Sea Salts/L) and plaque forming units (PFUs) enumerated using the agar overlay method (Sambrook and Russell 2001) with 3.5ml molten soft agar (20.5g Sigma Sea Salts and 6g low melting point agarose/L) and 300*µ*l overnight bacterial culture per plate.

### Community experiment

The community experiment was performed in triplicate with all five bacterial strains and all five phage strains together. Bacterial strains were inoculated and transferred individually; transfer cultures were pelleted at 4,000g for 10 minutes and 4 × 10^8^ cells of each strain (2 ×10^9^ total cells) were added to 200ml PC growth medium in each of 6x 2L flasks. Phage strains were then added at an MOI of 0.1 (4 ×10^7^ each phage, 2 × 10^8^ total phage) to three flasks; three contained no phage, and samples were taken every 35 minutes for 15 hours 45 minutes. At each time point, we sampled for OD_600_, CFUs, intracellular qPCR quantification (qINT), and extracellular qPCR quantification (qEXT).

Primers for qPCR were designed to amplify 75-150bp portions of each bacterial strain or phage, with negligible amplification from other members of the community. The primers were designed using the complete genomes of each bacterial strain or phage. All bacterial and phage genomes are sequenced and publicly available, but 2 of the bacterial genomes were not complete (*Cellulophaga baltica* 4 and *Pseudoalteromonas* sp. H100) and existed in genome fragments. To rectify this, we applied long-read sequencing to resequence and complete those two genomes. In addition, we also resequenced *Pseudoalteromonas* sp. 13-15 because primer design was difficult for the two *Pseudoalteromonas* bacterial strains as they are 99.9% identical genomically.

Primer pairs were tested for efficiency and mis-priming, and we used only primers with *>*85% efficiency and *<*10 copies/*µ*l amplification from other community members. DNA was extracted from qINT samples using the DNeasy Blood and Tissue kit (Qiagen) following the manufacturer’s instructions; qEXT samples were used as-is. qPCR was performed on an Eco Real-Time PCR System (Illumina) with PerfeCTa SYBR Green FastMix Reaction Mix (QuantaBio) in 13*µ*l reactions. Per reaction, we used 6.5*µ*l PerfeCTa master mix, 0.39*µ*l 10mM forward primer, 0.39*µ*l 10mM reverse primer (Supplementary Information Tab. S7), 4.72*µ*l nuclease-free water, and 1*µ*l template. For qINT samples, extracted DNA was used as template, and for qEXT, the 0.2*µ*m filtrate was used. Reactions were performed in technical duplicates with a standard curve consisting of 5x 10-fold dilution series of known concentration of the target strain or phage, used to calculate target sequence copies/*µ*l. Cycling conditions were as follows: polymerase activation for 5 minutes at 95^°^C; 40 cycles of 20 secs at 95^°^C, 10 secs at primer annealing temperature (Supplementary Information Tab. S7), and 20 secs at 72^°^C, and a 55^°^C – 95^°^C melt curve.

### Single bacteria and pairwise phage–bacteria experiments

Bacterial growth curves to determine doubling times were performed in triplicate in PC growth medium on cultures transferred 1:20 from overnight cultures into new media, grown stationary at RT. A regression of OD_600_ and colony-forming units (CFUs) was constructed using exponential phase time points to estimate cell density from OD readings.

Adsorption of phage to bacterial strains was characterized in triplicate by combining one phage with one bacterial strain and enumerating free (0.2*µ*m filtered to remove bacteria) and total phage over time. Pairs with a known interaction were combined at a multiplicity of infection (MOI) of 0.1 (10^7^ phage and 10^8^ cells/ml), and total and free phage PFUs were enumerated every 3-5 minutes for 24-25 minutes to calculate the adsorption constant. Pairs with no known interaction were combined at an MOI of 3 (3 × 10^8^ phage and 10^8^ cells/ml), and total and free phage PFUs were enumerated at time intervals for 3 hours to examine evidence in support of a statistically significant decrease in free phage. All infections were performed with strains in mid-exponential phase.

One-step growth curves were performed in triplicate to determine phage burst sizes and latent periods. Each phage–bacteria pair was combined at an MOI of 0.1 (10^7^ phage/ml and 10^8^ cells/ml) and incubated for 15 minutes to allow phage to adsorb to cells. The infection was then diluted 1:100 in PC growth medium to reduce the chances of new adsorptions. Total and free phage PFUs were enumerated at regular intervals for 2-4 hours. The conventional latent period was determined as the length of time before a significant increase in phage concentration. The average burst size is given by 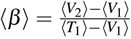 where *V* is the free phage density before (*V*_1_) and after (*V*_2_) the burst event, and *T*_1_ is the total phage density before the burst event. The averages ⟨·⟩ are taken over multiple time points and experiment replicates. All infections were performed with strains in mid-exponential phase.

### Pairwise experiment for multiple infection cycles

Each bacterial strain was inoculated, transferred, and pelleted as previously described. All pairwise phage-host interactions were tested in triplicate, with triplicate uninfected controls for each strain. Each strain was diluted to 2 × 10^6^ CFU/ml in PC media, and a single phage was added at an MOI of 0.1. From this infection, we immediately sampled to quantify total and free phage as previously described, and for intracellular (qINT) and extracellular (qEXT) qPCR. For qPCR, 1ml was pelleted at 10,000g for 5 minutes at 4^°^C. The supernatant was 0.2*µ*m filtered to remove any unpelleted bacteria and stored at 4^°^C for qEXT analysis and the pellet was flash frozen in liquid nitrogen and stored at -80^°^C for qINT analysis. We also aliquoted the infections into wells of a 96-well plate and read OD_600_ at 35-minute intervals for 15.75 hrs. At the end of this time, we repeated total phage, free phage, qEXT, and qINT sampling from the infection wells, pelleting 150*µ*l instead of 1ml. DNA was extracted from qINT pellets using the DNeasy Blood and Tissue kit (Qiagen), following the manufacturer’s instructions for Gram-negative bacteria. qPCR was performed to quantify intracellular and extracellular phage and bacterial DNA as previously described at 0, 3 hrs and 15.75 hrs.

### Modeling

We developed a coupled system of nonlinear ordinary differential equations (ODEs) to represent the dynamics of susceptible (*S*), exposed (*E*) and infected (*I*) bacterial cells and viruses (*V*). This SEIV model is parameterized by the viral latent period (*τ*), burst size (*β*), adsorption rate (*ϕ*), and bacterial growth rates (*r*). We utilized a non-stiff ODE solver with an adaptive time step^41^ and fixed constraints. Variation in the latent period distribution is incorporated through a compartmental model of *E* classes^17^ which generates Erlang distributed latent periods^42^. Supplementary Information Section S1.1 contains a full model description. The community SEIV model is scaled up to include interaction specific traits of all 9 phage-bacteria pairs. The SEIVD model also includes a state variable representing cellular debris (*D*), a proxy for the cumulative number of lysed cells. We assume debris inhibits infection via a Hill function given by 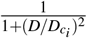 given the half-saturation constant 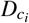. Supplementary Information Section S1.3 & S1.4 provides details on the community SEIV and SEIVD models, respectively.

### Inference

We used a delayed rejection adaptive Markov Chain Monte Carlo algorithm to fit the models to the data (including replicates when available) using Bayesian inference^43,44^. For the one-step growth experiments, the priors were derived from the conventional estimates of the life history traits. The priors and likelihood functions were chosen to be either truncated normal or log-normal distributions. For the community experiments over multiple infection cycles, the traits of the one-step experiment fail the prior predictive check. A coordinate gradient descent algorithm was used to fit one of the replicates^45^, which was used to inform priors, and the rest of the replicates were used to inform the likelihood in the Bayesian inference scheme. In all cases, posterior predictive checks and convergence tests were performed. Supplementary Information Section S1.6 provides a detailed description on inference methods.

## 3 Results

### Multiple bacteria and their associated phage coexist in the same community

We sought to assess whether a complex community of five bacteria that includes three *Cellulophaga baltica* (CBA) and two *Pseudoalteromonas* (PSA) isolated from similar environmental conditions could coexist at timescales spanning multiple life cycles when mixed with five associated virulent phage (see Methods in Sec. 2 for strain details). To do so, we first identified the host range of each of the five phage – identifying nine positive interactions among twenty-five combinations. The resulting infection matrix is modular such that the phage strains associated with either CBA or PSA do not cross-infect the other species (Fig. 1a). Next, we combined all five bacteria at the same initial densities and their five associated phage strains at a fixed multiplicity of infection (MOI) of 0.1. We sampled the *in vitro* community dynamics through triplicate experiments, recording cell and virion densities through intracellular quantitative Polymerase Chain Reaction (qPCR-INT) and extracellular quantitative Polymerase Chain Reaction (qPCR-EXT), respectively, every 35 minutes for 15.75 hrs, resulting in 28 time points for each flask (see Methods for further details on culture conditions and sampling methods). All 5 bacteria and 5 phage strains coexist for the duration of the experiment and the control (means and standard deviations shown in Fig. S1 and replicates shown in Fig. 1d). The high-resolution sampling of the phage–bacteria community also revealed subtle transient dynamics. Initially, phage populations decline given adsorption to cells even as cell populations increase given the relatively low initial impact of viral lysis. Subsequently, viral populations increase via infection and lysis (Fig. 1d-bottom row), leading to bacterial density declines ∼ 4–8 hrs after inoculation that subsequently stabilize to levels between 10^5^ cells/ml to 10^7^ cells/ml at the conclusion of the experiment. Similarly, phage populations stabilize between 10^9^ virions/ml to 10^11^ virions/ml, coexisting with, rather than eliminating, the bacterial populations. In contrast, a phage–free control experiment that otherwise followed the same protocols led to the coexistence of all five bacterial strains at densities between 10^7^ cells/ml to 10^9^ cells/ml (Fig. 1c). The total bacterial population in the phage-free community reached a value ∼ 10^9^ cells/ml than that with phage (Fig. 1b). We interpret these data as suggesting that the cumulative impacts of phage infection and lysis, and not resource scarcity, are limiting bacteria populations in the phage-treated community relative to that of the phage-free controls.

**Figure 1.**
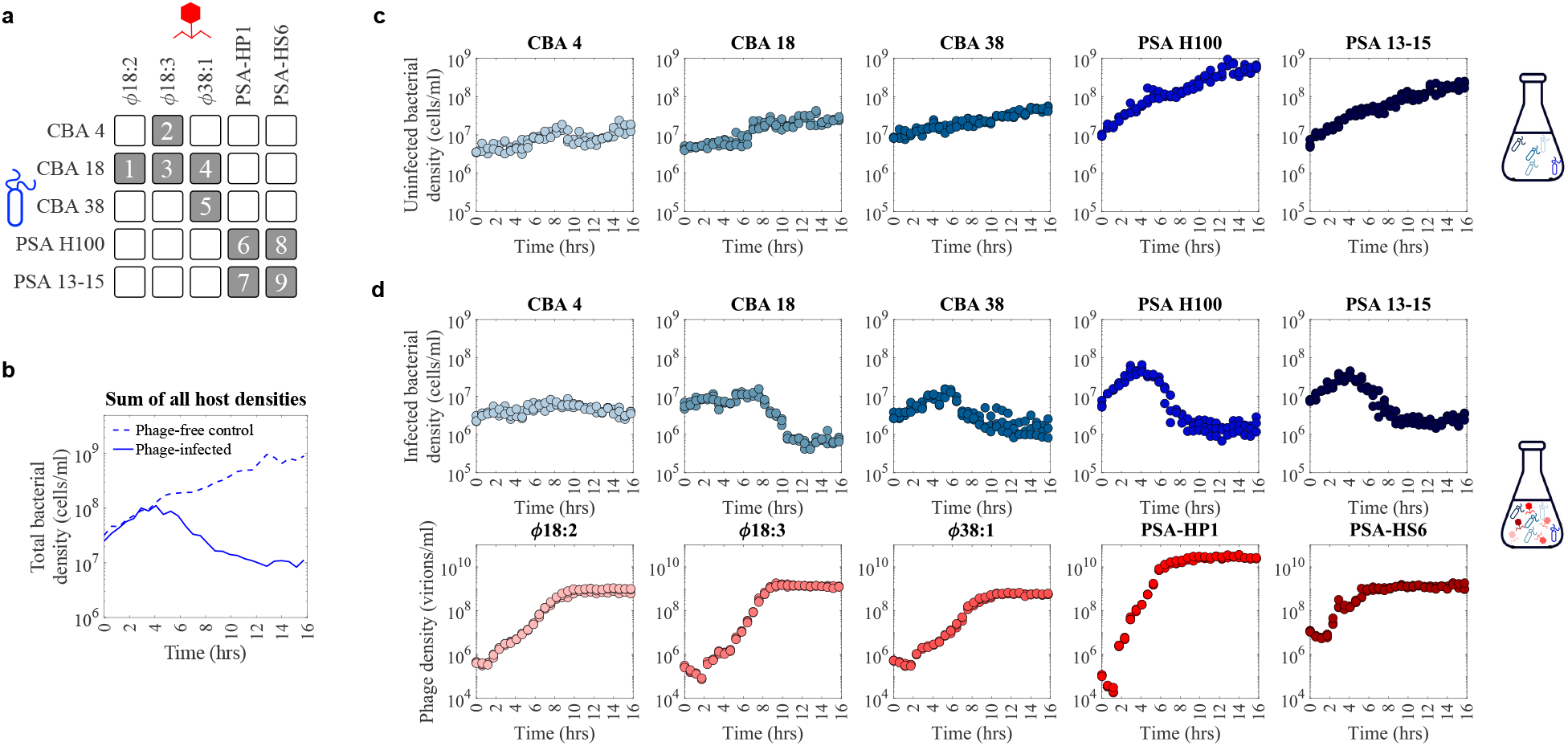
Community experimental data. **(a)** Adsorption assay experiments reveal an infection matrix with nine out of twenty-five possible infections occur among the strains, each numbered in a shaded gray box. Empty white boxes denote the absence of evidence for infection of the phage strain (in the columns) with the bacterial strain (in the rows). **(b)** Comparison of total bacterial population dynamics measured in triplicate via qPCR for the phage-free community control (five host strains) and the phage-infected community (five host and five phage strains). **(c)** Population dynamics of each host from the phage-free community control, where all the bacterial strains are grown together. **(d)** Population dynamics of each host and phage from the phage-infected community experiment. Five phage and five bacterial strains coexist together in the same flask. All three experimental replicates are shown for each panel. See Supplementary Information Fig. S1 for means and standard deviations of the population dynamics.

### Inferring pairwise phage–bacteria dynamics and traits

In order to reconcile the finding of elevated phage densities with community coexistence, we set out to quantify bacterial growth rates as well as intracellular and extracellular phage life history traits and then embed those into pairwise population models (Fig. 2a). We measured the growth rate *r* through doubling time experiments, finding that CBA strains divided every 4.2 – 5.8 hrs and PSA strains divided every 3.6 – 3.8 hrs (Fig. 2c). Next, we estimated the adsorption rates of each of the five phage to their respective hosts using adsorption assays with ranges between ∼ 2 × 10^−8^–2 × 10^−7^ ml/hr (Fig. 2c). New phage began to appear 0.5 hrs to 1.5 hrs later. For each interacting phage–bacteria pair, we used a Bayesian MCMC approach to fit a Susceptible-Exposed-Infected-Virus (SEIV) infection population model to one-step growth curves (Supplementary Information Fig. S2 and Methods S1.1, with prior distributions informed from conventional trait estimates). One-step growth experiments were not obtained for the interaction of phage *ϕ* 38:1 on CBA 18 due to methodical challenges associated with inefficient infections for this pair^46^. The SEIV model recapitulated phage production dynamics in one-step growth curves (Fig. 2b). Fitted models provide posterior estimates of life history traits including the adsorption rate *ϕ*, mean latent period *τ*, coefficient of variation of the latent period, CV of *τ*, and burst size *β* . We compared Bayesian inferences of traits with ‘conventional’ methods^15^ that utilize the time of first appearance of viruses and viral plateau to infer traits in Fig. 2c (as summarized in Supplementary Information Table S4). The conventional approach to inferring the mean latent period from one-step growth curves systematically underestimates the mean latent period by neglecting variability in latent period outcomes among infected cells in a population^17^. We estimate the coefficient of variation of the latent period to be ≈ 0.1 for most strains. As is shown in Fig. 2b, the nonlinear SEIV model of phage-bacteria interactions with inferred life history traits can recapitulate virus production dynamics over a single lytic period.

**Figure 2.**
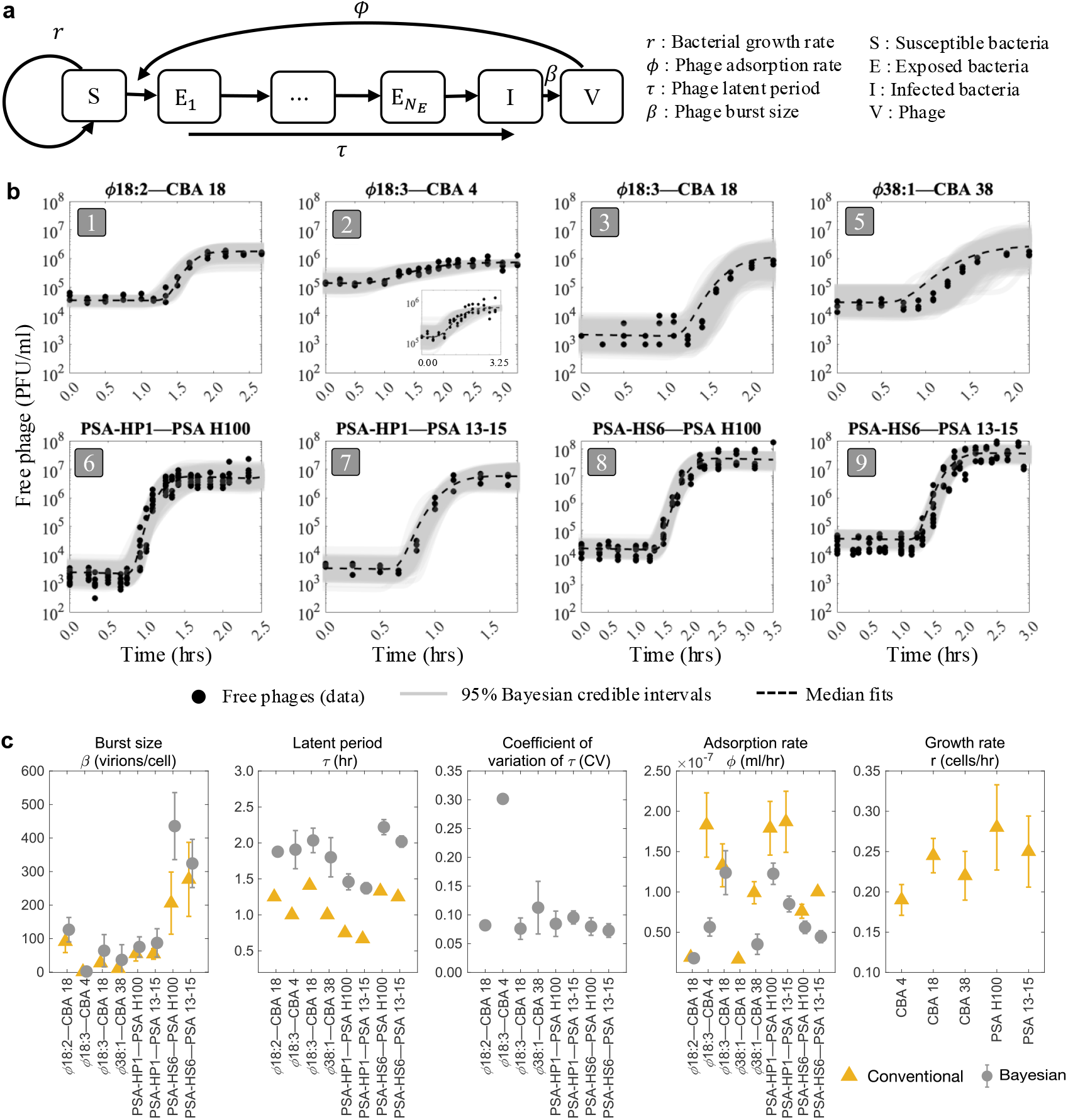
Pairwise interactions between phage and bacteria. **(a)** Pairwise phage-bacteria dynamics fit with a nonlinear ODE model (described in Supplementary Information S1.1) with a variable number of compartmental stages of infection *N*_*E*_ (Supplementary Information Fig. S2), parameterized by the specific phage–bacteria pairwise life history traits: *r, ϕ, τ, β* . **(b)** Free phage density measured through one-step growth curves (black dots) for eight out of nine interacting phage–bacteria pairs. Using the model in (a), the pairwise life history parameters *β, τ*, and *ϕ* are inferred by a Bayesian algorithm. The Bayesian fits are shown in gray lines along with 95% credible interval shaded for each of them. **(c)** The conventionally inferred life history traits from single-strain experiments (yellow triangles) such as one-step growth curves, pairwise adsorption assays, and bacterial growth doubling time experiments are compared to Bayesian inferred life history traits (gray circles) from one-step growth curves; standard deviations shown as error bars. The coefficient of variation for the latent period distribution (related to the inferred number of compartments *N*_*E*_ as 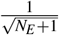) is also inferred for each pair. Full details on the fitting statistics are provided in Supplementary Information Fig. S9-S10.

### Dynamics of pairwise infections over multiple infection cycles

As a step towards understanding the basis for coexistence at the community scale, we set out to understand if phage-bacteria pairs could persist together over multiple infection cycles. We reinitiated all 8 viable, pairwise infection experiments focusing on phage-bacteria pairs for which we have robust parameter estimates (as described in the prior section). In each case, we measured host densities via qPCR at *t* = 0, *t* = 3.5 hrs (the longest endpoint within the trait inference experiments) and *t* = 15.75 hrs (consistent with time scales in the community experiment; see Sec. 2). The ratio of final to initial host density in control (phage-free) populations varied from 10 to nearly 100, consistent with host proliferation in the absence of phage (Fig. 3). We hypothesized that addition of phage would lead to the collapse of bacterial populations. Instead, host populations persisted in all pairwise infection cases to the end of the experiment. In all cases, the ratio of final to initial host density in infected populations was lower than that in control populations (see Fig. 3 and Fig. S3 for further details).

**Figure 3.**
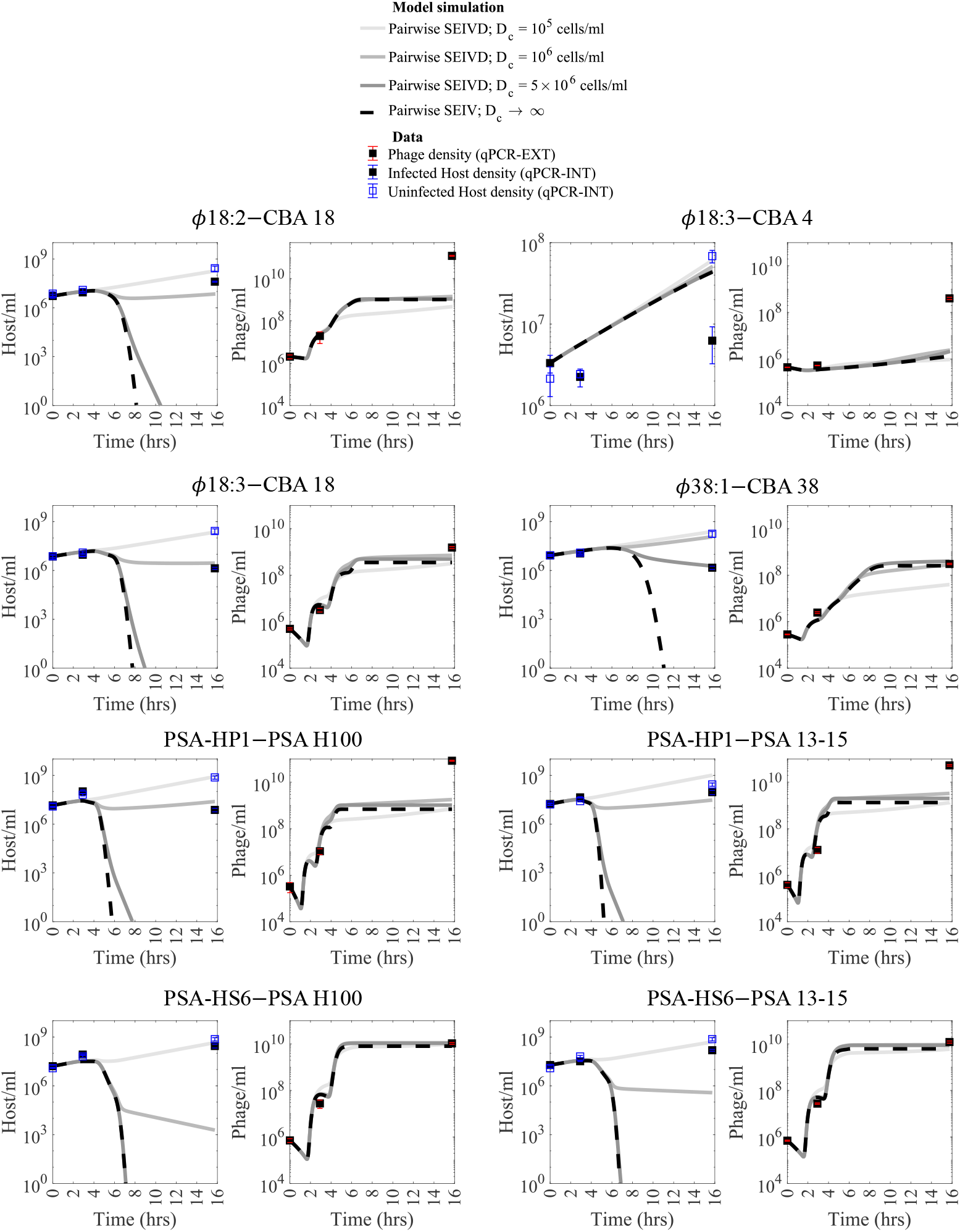
Host densities from pairwise infection experiments over multiple life cycles. For 8 phage-host infection pairs, phage and host densities obtained via triplicate qPCR-INT (intracellular) and qPCR-EXT (extracellular) measurements respectively for 15.75 hrs (representing multiple life cycles) are plotted in solid squares. Phage-free control for hosts are plotted in open squares. Pairwise SEIV model (dashed lines; where *D*_*c*_ → ∞ and there is no debris-mediated attenuation of infection) and SEIVD model (solid lines) parameterized through one-step experiments (Fig. 2) and experimentally recorded initial densities are plotted for each experimental pair. Pairwise SEIV model predicts host elimination in 7 out of the 8 cases in these pairwise experiments, whereas pairwise SEIVD model with infection attenuation qualitatively demonstrates phage-host coexistence for experimental timescales. For the ratios of final to initial host densities please refer to Fig. S3.

We simulated pairwise SEIV models over multiple infection cycles to see if we could recapitulate the pairwise experimental findings, assuming life history traits inferred via single-cycle pairwise experiments, as shown in the prior section. The pairwise SEIV models predicted the experimentally observed densities up to 3 hrs, but then failed to predict densities over longer time scales. Instead of persistence, the pairwise SEIV models predict population crashes within 16 hrs for 7 out of the 8 host phage pairs (dashed lines in Fig. 3) and predict substantial host population declines of at least a factor of 10^2^ in all cases, except for CBA 4 interacting with *ϕ* 18 : 3. In contrast, we found that 5 out of 8 experiments with infected populations exhibited increased host densities and never exhibited host population declines greater than a factor of 10^2^. This finding implies that host growth remains largely in balance and potentially exceeds viral lysis in the majority of pairs, in strong contrast to simulated dynamics in models that inferred apparently efficient lysis within one-step growth experiments.

We hypothesized that the pairwise SEIV models neglected a density-dependent feedback that was not observed when fitting to growth-dominated early time data. As such, we extended the SEIV model to include the impact of a negative feedback mechanism in which bacterial lysis inhibits new infections, e.g., potentially due to inhibitory effects of cellular debris on viral adsorption and lysis. Rather than representing a confirmed mechanistic pathway, the SEIVD model is an effective description of cellular-level mechanisms that could impede phage infection centered around one phenomenological feature: infection attenuation increases nonlinearly as a function of lysed cells. This new pairwise SEIVD model (where ‘D’ represents debris, a proxy for lysed cells, see Model S1.2) includes a negative feedback loop in which the strength of infection attenuation is given by a Hill function tuned by a critical concentration, *D*_*c*_. We choose a Hill coefficient of 2 for all cases. Given *D*_*c*_ values ranging from 10^5^ to 5 × 10^6^ cells/ml, the model predicts bacterial persistence with minimal impacts on phage population dynamics. This finding suggests that additional stabilizing mechanisms can be present even in pairwise contexts at high densities that may also be relevant when analyzing community dynamics.

### Scaling-up pairwise infection models do not recapitulate community level dynamics

We scaled up the pairwise SEIV and SEIVD models to a community context by incorporating cross-infections as obtained through adsorption assay experiments (Fig. 1a). In a community model each susceptible bacteria population, *i*, with density *S*_*i*_ can be infected by each of its associated phage, *j*, with density *V*_*j*_. For each of the nine pairs, we account for latent period heterogeneity by modulating the number of classes of ‘exposed’ compartments, *E*_*i j*_. We specifically excluded the potential for multiple infections of a single cell, either by the same or different phage. The community models include the potential to re-estimate life history traits and a strain-specific critical debris concentration, 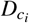 (Fig. 4). Complete details of the community SEIV and SEIVD models are described in Supplementary Information S1.3-S1.4.

**Figure 4.**
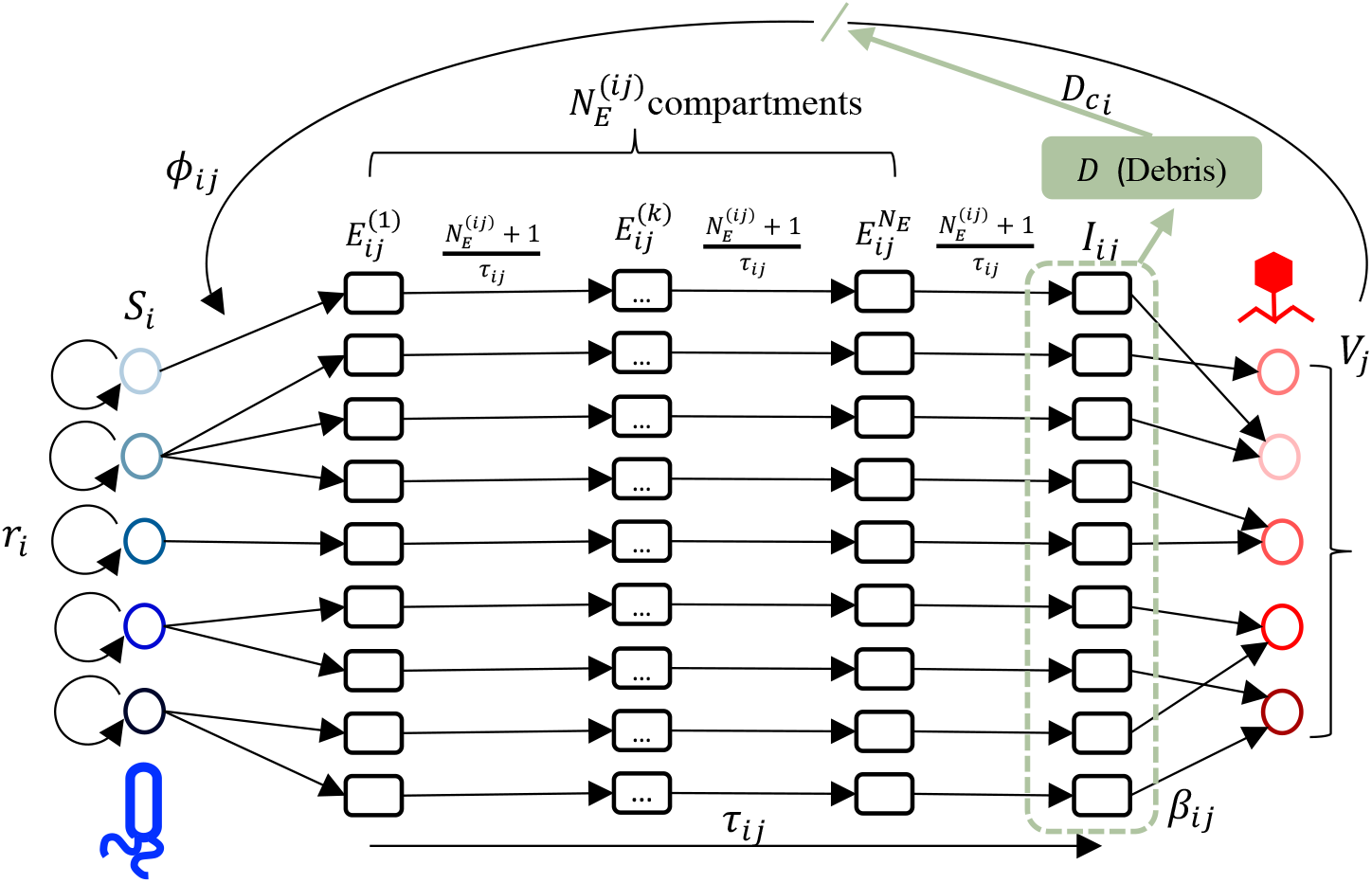
SEIV and SEIVD based scaled-up community model. Using the infection matrix in Fig. 1(b) the model is scaled up to include the 9 interactions between 5 bacterial and 5 phage strains and respective number of exposed classes 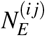 for each interaction. *S*_*i*_, *E*_*i j*_ and *I*_*i j*_ are susceptible, exposed, and infected stages in the sequential infection for each bacteria *i* by phage *j*. This SEIV community model is parameterized by the phage infection traits of the constituent pairs of the community, such as latent periods (*τ*_*i j*_), adsorption rates (*ϕ*_*i j*_), burst sizes (*β*_*i j*_) and bacterial growth rates (*r*_*i*_). For the SEIVD model, we include a state variable *D*, which represents the dead cell density (green box) that can attenuate new infections, given a critical debris concentration 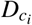. See main text and Supplementary Information S1.3 and S1.4 for the full description of the SEIV and SEIVD models, respectively.

We simulated the community SEIV model using the pairwise Bayesian median posterior estimates of the viral burst sizes, adsorption rates, average latent periods and their variability for the eight inferred interactions, and bacterial growth rates (Fig. 2c and Supplementary Information Table S4, and Fig. S9-S10). As noted, one-step phage growth parameters could not be inferred for the infection of *ϕ* 38:1 on CBA 18. Instead, we used a range of parameter values derived from the literature^46^. The community SEIV model parameterized by pairwise-fitted infection traits failed to capture the observed community dynamics. As in the pairwise simulations over multiple infection cycles (Fig. 3), the community SEIV model predicts that all bacterial populations should decrease rapidly given the explosive growth of phage populations (Fig. 5a). Even when accounting for variability in life history traits, the models predict decreases exceeding two orders of magnitude within 10 hrs post-exposure to viruses. This bacterial population collapse (not observed in experiments and akin to that shown in pairwise SEIV models) arises via rapid increases and then stabilization of viral populations.

**Figure 5.**
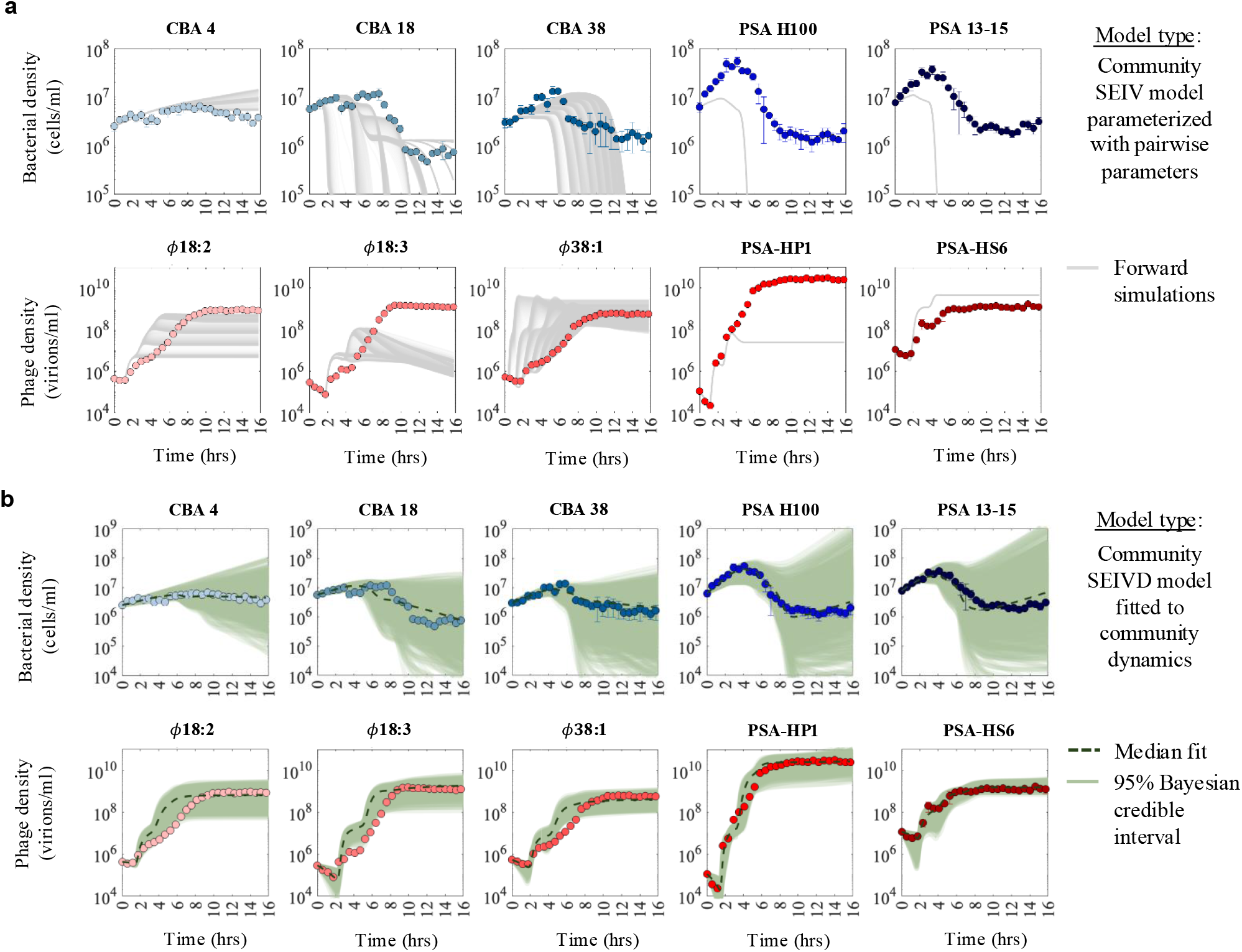
Comparison of observed versus simulated phage–bacteria population dynamics. The mean of the population density time series for all the five bacterial hosts and five phage obtained from triplicate experiments are shown (circles), with their respective standard deviations. **(a)** The life history traits inferred from single strain experiments and one-step growth curves in Fig. 2 are directly used to parameterize the scaled-up SEIV community model, where uncertainty in model outcome (gray lines) is associated with uncertainty regarding the burst size and latent period of *ϕ* 38:1 on CBA 18 derived from literature (parameters summarized in Supplementary Information Table S4). **(b)** The community SEIVD model is used to infer community-level life history traits. Simulations from 95% Bayesian credible intervals of the posterior distribution are plotted in green and median fits in dashed lines. Priors and posterior distributions are detailed in Supplementary Information Fig. S12. Additional information on fits is available in Supplementary Information Fig. S13 for the MCMC chains and Supplementary Information Fig. S14-S15 for the MCMC convergence statistics.

### Integrating higher-order interactions and density-dependent feedback in the community model enables complex coexistence

We hypothesize that density-dependent feedback can stabilize complex phage-host communities. To evaluate this hypothesis, we simulated the SEIVD community model parameterized by pairwise parameters and a broad range of independently sampled 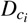 values utilized in simulating the pairwise experiments. We observed that the community model with density-dependent infection attenuation prevents extinction of host strains but fails to quantitatively recapitulate the phage or host densities (Fig. S4a). One potential explanation is that life history traits of phage might differ in community vs. pairwise contexts. To evaluate this hypothesis, we refit the community SEIV model to the first 6.4 hrs of community dynamics, intentionally excluding density-dependent infection attenuation (see Supplementary Information Section S1.6.4). The re-fit community SEIV model quantitatively captured the population dynamics of all phage strains for the entire 15.75 hr experiment (Fig. S4b-bottom) while only capturing bacterial dynamics for the first ∼ 6 hrs (Fig. S4b-top). Nevertheless, the improvement of community SEIV model fits on the shorter timescale suggests that viral life history traits differ between pairwise and community contexts (Supplementary Information Tab. S9).

Finally, we combined both putative stabilizing mechanisms together in a single model: refitting the SEIVD model by performing MCMC-based Bayesian inference on the triplicate population time series data to estimate the joint posterior distribution of the phage–bacteria life history traits and 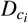 values. This SEIVD community model can quantitatively fit both the experimentally recorded bacterial and phage time series data, unlike either the scaled-up pairwise model with density-dependent infection attenuation or the re-fit SEIV community model (Fig. 5b, additional robustness analysis in Fig S5 and S6, parameter summary and statistics in Table S5 and Fig. S12-S15). The model-predicted phage dynamics include both initial decreases associated with virion adsorption and subsequent production that eventually stabilizes at levels consistent with observations. Likewise, the ensemble of bacterial density dynamics includes stabilization in all cases, as well as the potential for bacterial population rebounds even in the absence of phage-resistance.

### Experimentally testing putative higher-order interaction mechanisms

We set out to test both of the hypothesized mechanisms that shape community-scale coexistence in the SEIVD model: higher order interactions and density-dependent infection attenuation. First, we compared viral traits inferred from pairwise experiments and community experiments inferred via the pairwise SEIV and community SEIVD. We found moderate to strong shifts in traits for 11 trait parameters, whereas 8 of them showed minor differences, and 5 were inconclusive (Fig 6 and Table S6 for statistical comparison). The SEIVD community model predicted strong shifts in burst sizes, e.g., the community estimated burst size of PSA-HP1 is larger and that of PSA-HS6 is smaller compared to values inferred in pairwise experiments. To test these unexpected shifts, we performed one-step growth experiments of the PSA strains with all hosts combined (1 phage with the 5 host subcommunity). We found that in the subcommunity of PSA hosts the burst size of PSA-HP1 increases, whereas the burst size of PSA-HS6 decreases relative to the pairwise case, consistent with model-inferred predictions (Supplementary Information S2.3, Fig. S7).

**Figure 6.**
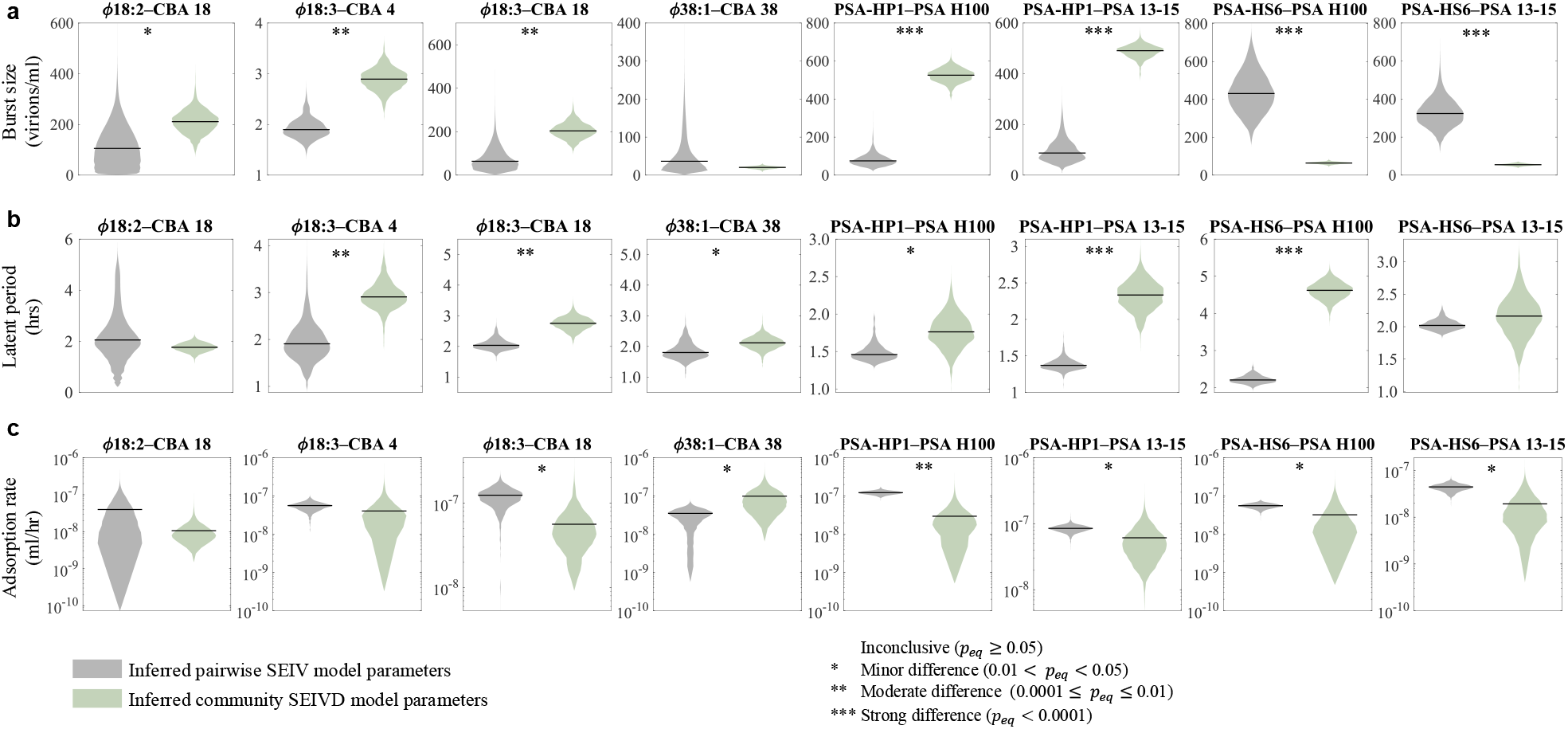
Violin plots for pairwise versus community phage traits. Phage traits inferred via Bayesian methods with pairwise SEIV and community SEIVD models (corresponding to the fitted dynamics in Fig. 2b and Fig. 5b respectively) are compared with violin plots for the 8 phage-host interactions described in Fig. 2. Sub-figures a-c show burst size, latent period and adsorption rate respectively. We compared the traits using a Bayesian test for equivalence^70^ and the posterior probability of practical equivalence, *p*_*eq*_, is defined through a pre-specified region of equivalence (analogous to P value). Methods are outlined in Supplementary Information section S1.7, see Supplementary Information Table S6 for detailed statistics.

In order to test for density-dependent attenuation of infection, we added bacteria-filtered and phage-inactivated spent media from 12 hrs of the community experiment to fresh pairwise infections for all 9 phage-bacteria infection pairs. In each case, we contrasted dynamics with that of including the host alone and the phage-bacteria infection pair without community spent media. Host densities remained elevated for all strains in the phage-free control. For 3 CBA infections and 1 PSA infection, namely, *ϕ* 18 : 3 on CBA 4, *ϕ* 18 : 3 on CBA 18, *ϕ* 38 : 1 on CBA 38, and PSA-HP1 on PSA 13-15, the host densities fall below the limit of detection in the infection-only control (without community lysate). In contrast, the host persists when community spent media is added along with the phage (Supplementary Information Fig. S8), consistent with a density-dependent attenuation of infection and lysis.

## 4 Discussion

In this work, we explored mechanisms of coexistence within a synthetic marine phage–bacteria community. In contrast to model predictions derived from short-term pairwise phage-bacteria experiments, we found that individual phage-bacteria pairs and complex multi-strain communities can coexist over multiple infection cycles. Through a process of iterative model-data integration, we hypothesized that higher-order interactions stabilize community dynamics. These interactions include context-dependent shifts in life history traits and infection attenuation at relatively high viral densities. By incorporating both types of higher-order interaction mechanisms, we quantitatively recapitulate community time series data and the coexistence of all 5 heterotrophic bacteria and 5 associated phage over a 15.75 hr time series experiment (far longer than the typical infection cycles of 1-2 hrs). In doing so, we also experimentally validated the model-inferred higher-order interaction mechanisms: (i) higher order interactions such that life history traits exhibit directional shifts in community vs. pairwise experiments, and, (ii) density-dependent attenuation, such that phage lysates diminish the intensity of infection-induced lysis.

Central to our analysis was the development of a nonlinear population dynamics model that incorporated the combined effects of density-dependent infection attenuation and higher order interactions. Both mechanisms were essential to reconcile the potential for rapid lysis when viral densities are low and the coexistence of bacteria with viruses when viral densities are high. First, our finding of density-dependent infection attenuation might be due to accumulation of cellular debris that changes host physiology and/or viral infectiousness^47^. The spent media experiments alone (Fig. S8) do not necessarily rule out additional inhibitory effects that might occur via feedback in a growing bacterial community. This finding also suggests caution is warranted in extrapolating from short-to long-term dynamics. Ecological models that are fit to dynamics in experimentally convenient conditions may not yet fully represent feedback across biologically relevant regimes. Second, higher-order interactions are increasingly considered to be potentially powerful drivers of coexistence^19,48,49^, including in microbial communities^24,25,50–54^. Our work reinforces the ecological relevance of earlier findings that life history traits can also depend on context^26,55^, e.g., phage latent period may decrease in response to lower resources^56^ and phage growth can be affected by bacterial growth rate^57^.

Despite being able to quantitatively recapitulate complex community dynamics, our modeling and inference framework comes with caveats. A salient feature of our ecological models is the explicit integration of the variability in the latent period observed in prior phage host systems^58–61^ modeled via multi-compartment stages of infection progression^17,62^. For simplicity, we assumed that the variability in this latent period is the same for all pairwise interactions in the community SEIVD model. In reality, we expect variability to be strain dependent. Another key assumption is that bacterial traits are constant in time, whereas long-term coexistence would need to account for nuanced bacteria-bacteria interactions as well. Further, we have assumed that each bacterial strain is infected by at most one virus. Previous studies have shown that the outcomes of infection may differ depending on the multiplicity of infection^63–65^ and/or communication systems^66^. Additionally, in the community and the pairwise SEIVD models, infection attenuation only becomes relevant at later stages of the experiments. Multiple biochemical pathways that might be responsible for infection attenuation at timescales of multiple infection cycles or at higher cell densities, including those mediated via debris, is abstracted through this effective mechanism. Finally, we have not included the impacts of phage-resistance or counter-defense amongst phage – nor did we observe the emergence of phage-resistant bacterial strains in our experiments. The repeatability of strain dynamics and the absence of bacterial regrowth even as total densities were 100-fold less than observed in phage-free controls suggests that phage-resistance (even if present) is not ecologically relevant on these time scales in this system. Exploring the interface between resistance and counter-resistance will be critical in understanding longer term, eco-evolutionary dynamics of phage-bacteria communities.

In summary, our work suggests that pairwise interactions inform but do not wholly determine community-level interactions. We find that higher-order interactions matter and that assuming rapid pairwise lysis does not necessarily explain phage–bacteria community dynamics and coexistence. The community models and inference framework developed here may be of service in exploring the generality of these feedback mechanisms in environmental, agricultural, and human-associated microbiomes^67–69^. In doing so, our work also presents a caution: scaling up results from pairwise interaction outcomes to community dynamics will likely require accounting for density-dependent feedback and higher-order interactions. Identifying the relevance of both mechanisms is likely to improve the predictive capacity of community-scale models as well as provide into the coexistence of diverse populations of viruses and bacteria.

## Data availability

The datasets and code used for this project are available at: https://github.com/RaunakDey/VIMIMO paper and are archived on Zenodo, https://doi.org/10.5281/zenodo.17526176.

## Author contributions

MBS and JSW conceived the study. RD and ARC led theory and analyses with support from SJB, DD, DM, and JSW. NS led experiment design/implementation and data acquisition with support from ARC, CLS, MB, AM, and JG. CLS led sequencing and primer design. RD, ARC, and JSW wrote the manuscript with support from NS, DD, CLS, SJB, and MBS. RD and ARC generated tables and figures.

## Acknowledgments

We thank Steven Wilhelm, Gary LeCleir, Naomi Gilbert, and Debbie Lindell for their thoughtful feedback. We also thank members from the Weitz Group, Gabi Steinbach and Jessica Irons for administrative support and Tapan Goel for code review. We thank members of the Sullivan group Shaun Maxwell, Yueh-Fen Li, Guillermo Dominguez-Huerta, Lauren Chittick, and Siddharth Bhindwallam for help with preparing/sampling the synthetic community. The authors acknowledge the University of Maryland supercomputing resources (http://hpcc.umd.edu), Partnership for an Advanced Computing Environment (PACE) at the Georgia Institute of Technology, and Ohio Supercomputer Center made available for conducting the research reported in this paper. This work was supported by the National Science Foundation under Grant #1829636 (J.S.W.), Simons Foundation grant #72123 (J.S.W.), NSF EMERGE Biology Integration Institute award #2022070 (M.B.S.), and DOE Systems Biology award #DE-SC0023307 (M.B.S.). R.D. acknowledges fellowship support from the University of Maryland Center of Excellence in Microbiome Sciences and the Thomas G. Mason Interdisciplinary Physics Fund. S.J.B. and J.S.W. are investigators at the University of Maryland-Institute for Health Computing, which is supported by funding from Montgomery County, Maryland and The University of Maryland Strategic Partnership: MPowering the State, a formal collaboration between the University of Maryland, College Park and the University of Maryland, Baltimore.

## References

1. Chevallereau, A., Pons, B. J., van Houte, S. et al. Interactions between bacterial and phage communities in natural environments. Nat. Rev. Microbiol. 20, 49–62 (2022).

2. Mirzaei, M. K. & Maurice, C. F. Ménage à trois in the human gut: interactions between host, bacteria and phages. Nat. Rev. Microbiol. 15, 397–408 (2017).

3. Salmond, G. P. C. & Fineran, P. C. A century of the phage: past, present and future. Nat. Rev. Microbiol. 13, 777–786 (2015).

4. Suttle, C. A. Marine viruses —major players in the global ecosystem. Nat. Rev. Microbiol. 5, 801–812 (2007).

5. Rohwer, F. & Thurber, R. V. Viruses manipulate the marine environment. Nature 459, 207–212 (2009).

6. Weitz, J. S. & Wilhelm, S. W. Ocean viruses and their effects on microbial communities and biogeochemical cycles. F1000 biology reports 4, 17–17 (2012).

7. Flores, C. O., Meyer, J. R., Valverde, S. et al. Statistical structure of host–phage interactions. Proc. Natl. Acad. Sci. 108, E288 (2011).

8. Flores, C. O., Valverde, S. & Weitz, J. S. Multi-scale structure and geographic drivers of cross-infection within marine bacteria and phages. The ISME J. 7, 520–532 (2013).

9. Weitz, J. S. Quantitative viral ecology: Dynamics of viruses and their microbial hosts. Princet. Press. (2016).

10. Kauffman, K. M., Chang, W. K., Brown, J. M. et al. Resolving the structure of phage–bacteria interactions in the context of natural diversity. Nat. Commun. 13, 372 (2022).

11. Breitbart, M., Bonnain, C., Malki, K. et al. Phage puppet masters of the marine microbial realm. Nat. Microbiol. 3, 754–766 (2018).

12. Weitz, J. S., Poisot, T., Meyer, J. R. et al. Phage–bacteria infection networks. Trends Microbiol. 21, 82–91 (2013).

13. Borin, J. M., Lee, J. J., Lucia-Sanz, A. et al. Rapid bacteria-phage coevolution drives the emergence of multiscale networks. Science 382, 674–678 (2023).

14. Layeghifard, M., Hwang, D. M. & Guttman, D. S. Disentangling interactions in the microbiome: A network perspective. Trends Microbiol. 25, 217–228 (2017).

15. Abedon, S. T. Advances in molecular and cellular microbiology. Camb. Univ. Press. (2008).

16. Hyman, P. & Abedon, S. T. Practical methods for determining phage growth parameters. Bacteriophages: methods protocols, volume 1: isolation, characterization, interactions 175–202 (2009).

17. Dominguez-Mirazo, M., Harris, J. D., Demory, D. et al. Accounting for cellular-level variation in lysis: implications for virus–host dynamics. mBio 15, e01376–24 (2024).

18. Nemergut, D. R., Schmidt, S. K., Fukami, T. et al. Patterns and processes of microbial community assembly. Microbiol. Mol. Biol. Rev. 77, 342–356 (2013).

19. Friedman, J., Higgins, L. M. & Gore, J. Community structure follows simple assembly rules in microbial microcosms. Nat. Ecol. & Evol. 1, 109 (2017).

20. Goldford, J. E., Lu, N., Bajić, D. et al. Emergent simplicity in microbial community assembly. Science 361, 469–474 (2018).

21. Ratzke, C., Barrere, J. & Gore, J. Strength of species interactions determines biodiversity and stability in microbial communities. Nat. Ecol. & Evol. 4, 376–383 (2020).

22. Grilli, J., Barabás, G., Michalska-Smith, M. J. et al. Higher-order interactions stabilize dynamics in competitive network models. Nature 548, 210–213 (2017).

23. Bairey, E., Kelsic, E. D. & Kishony, R. High-order species interactions shape ecosystem diversity. Nat. Commun. 7, 12285 (2016).

24. Sanchez-Gorostiaga, A., Bajić, D., Osborne, M. L. et al. High-order interactions distort the functional landscape of microbial consortia. PLoS Biol. 17, e3000550 (2019).

25. Chang, C.-Y., Bajić, D., Vila, J. C. et al. Emergent coexistence in multispecies microbial communities. Science 381, 343–348 (2023).

26. van den Berg, N. I., Machado, D., Santos, S. et al. Ecological modelling approaches for predicting emergent properties in microbial communities. Nat. Ecol. & Evol. 6, 855–865 (2022).

27. Chesson, P. Mechanisms of maintenance of species diversity. Annu. Rev. Ecol. Syst. 31, 343–366 (2000).

28. Ellner, S. P., Snyder, R. E., Adler, P. B. et al. An expanded modern coexistence theory for empirical applications. Ecol. Lett. 22, 3–18 (2019).

29. Lear, K. O., Whitney, N. M., Morris, J. J. et al. Temporal niche partitioning as a novel mechanism promoting co-existence of sympatric predators in marine systems. Proc. Royal Soc. B 288, 20210816 (2021).

30. Rivett, D. W. & Bell, T. Abundance determines the functional role of bacterial phylotypes in complex communities. Nat. Microbiol. 3, 767–772 (2018).

31. Tikhonov, M. Community-level cohesion without cooperation. Elife 5, e15747 (2016).

32. Koskella, B. & Brockhurst, M. A. Bacteria–phage coevolution as a driver of ecological and evolutionary processes in microbial communities. FEMS Microbiol. Rev. 38, 916–931 (2014).

33. Fortuna, M. A., Barbour, M. A., Zaman, L. et al. Coevolutionary dynamics shape the structure of bacteria-phage infection networks. Evolution 73, 1001–1011 (2019).

34. Blazanin, M. & Turner, P. E. Community context matters for bacteria-phage ecology and evolution. The ISME J. 15, 3119–3128 (2021).

35. Holmfeldt, K., Solonenko, N., Shah, M. et al. Twelve previously unknown phage genera are ubiquitous in global oceans. Proc. Natl. Acad. Sci. 110, 12798–12803 (2013).

36. Duhaime, M. B., Solonenko, N., Roux, S. et al. Comparative omics and trait analyses of marine Pseudoalteromonas phages advance the phage OTU concept. Front. Microbiol. 8, 1241 (2017).

37. Deng, L., Gregory, A., Yilmaz, S. et al. Contrasting life strategies of viruses that infect photo- and heterotrophic bacteria, as revealed by viral tagging. mBio 3, 10.1128/mbio.00373–12 (2012).

38. Holmfeldt, K., Middelboe, M., Nybroe, O. et al. Large variabilities in host strain susceptibility and phage host range govern interactions between lytic marine phages and their Flavobacterium hosts. Appl. Environ. Microbiol. 73, 6730–6739 (2007).

39. Wichels, A., Biel, S. S., Gelderblom, H. R. et al. Bacteriophage diversity in the North Sea. Appl. Environ. Microbiol. 64, 4128–4133 (1998).

40. Melissa, D., Antje, W. & Matthew, S. Six Pseudoalteromonas strains isolated from surface waters of Kabeltonne, Offshore Helgoland, North Sea. Genome Announc. 4, e01697–15 (2016).

41. Shampine, L. F. & Reichelt, M. W. The MATLAB ODE suite. SIAM J. on Sci. Comput. 18, 1–22 (1997).

42. Hurtado, P. J. & Kirosingh, A. S. Generalizations of the ‘Linear Chain Trick’: incorporating more flexible dwell time distributions into mean field ode models. J. Math. Biol. 79, 1831–1883 (2019).

43. Haario, H., Saksman, E. & Tamminen, J. An adaptive metropolis algorithm. Bernoulli 223–242 (2001).

44. Haario, H., Laine, M., Mira, A. et al. DRAM: efficient adaptive MCMC. Stat. Comput. 16, 339–354 (2006).

45. Wright, S. J. Coordinate descent algorithms. Math. Program. 151, 3–34 (2015).

46. Dang, V. T., Howard-Varona, C., Schwenck, S. et al. Variably lytic infection dynamics of large Bacteroidetes podovirus phi38:1 against two Cellulophaga baltica host strains. Environ. Microbiol. 17, 4659–4671 (2015).

47. Talmy, D., Beckett, S. J., Zhang, A. B. et al. Contrasting controls on microzooplankton grazing and viral infection of microbial prey. Front. Mar. Sci. 6, 182 (2019).

48. Mayfield, M. M. & Stouffer, D. B. Higher-order interactions capture unexplained complexity in diverse communities. Nat. Ecol. & Evol. 1, 0062 (2017).

49. Billick, I. & Case, T. J. Higher order interactions in ecological communities: what are they and how can they be detected? Ecology 75, 1529–1543 (1994).

50. Mickalide, H. & Kuehn, S. Higher-order interaction between species inhibits bacterial invasion of a phototroph-predator microbial community. Cell Syst. 9, 521–533 (2019).

51. Datta, M. S., Sliwerska, E., Gore, J. et al. Microbial interactions lead to rapid micro-scale successions on model marine particles. Nat. Commun. 7, 11965 (2016).

52. Hu, J., Amor, D. R., Barbier, M. et al. Emergent phases of ecological diversity and dynamics mapped in microcosms. Science 378, 85–89 (2022).

53. Foster, K. R. & Bell, T. Competition, not cooperation, dominates interactions among culturable microbial species. Curr. Biol. 22, 1845–1850 (2012).

54. Sanchez-Martinez, R., Arani, A., Krupovic, M. et al. Episomal virus maintenance enables bacterial population recovery from infection and promotes virus-bacterial coexistence. The ISME J. wraf066 (2025).

55. Schmitz, O. J., Buchkowski, R. W., Burghardt, K. T. et al. Functional traits and trait-mediated interactions: connecting community-level interactions with ecosystem functioning. Adv. Ecol. Res. 52, 319–343 (2015).

56. Abedon, S. T., Herschler, T. D. & Stopar, D. Bacteriophage latent-period evolution as a response to resource availability. Appl. Environ. Microbiol. 67, 4233–4241 (2001).

57. Nabergoj, D., Modic, P. & Podgornik, A. Effect of bacterial growth rate on bacteriophage population growth rate. Microbiol. Open 7, e00558 (2018).

58. Doermann, A. H. Lysis and lysis inhibition with Escherichia Coli bacteriophage. J. Bacteriol. 55, 257–276 (1948).

59. Dennehy, J. J. & Wang, I.-N. Factors influencing lysis time stochasticity in bacteriophage λ . BMC Microbiol. 11, 1–12 (2011).

60. Wedd, C., Yunusov, T., Smith, A. et al. Single-cell imaging of the lytic phage life cycle in bacteria. bioRxiv 2024–04, DOI: 10.1101/2024.04.11.588870v4 (2024).

61. Baker, C. W., Miller, C. R., Thaweethai, T. et al. Genetically determined variation in lysis time variance in the bacteriophage ϕx174. G3: Genes, Genomes, Genet. 6, 939–955 (2016).

62. Hinson, A., Papoulis, S., Fiet, L. et al. A model of algal-virus population dynamics reveals underlying controls on material transfer. Limnol. Oceanogr. 68, 165–180 (2023).

63. Bondy-Denomy, J., Qian, J., Westra, E. R. et al. Prophages mediate defense against phage infection through diverse mechanisms. The ISME J. 10, 2854–2866 (2016).

64. Mavrich, T. N., Hatfull, G. F., Hiller, N. L. et al. Evolution of superinfection immunity in cluster a mycobacteriophages. mBio 10, e00971–19 (2021).

65. Berngruber, T. W., Weissing, F. J. & Gandon, S. Inhibition of superinfection and the evolution of viral latency. J. Virol. 84, 10200–10208 (2010).

66. Erez, Z., Steinberger-Levy, I., Shamir, M. et al. Communication between viruses guides lysis–lysogeny decisions. Nature 541, 488–493 (2017).

67. Geesink, P., Ter Horst, J. & Ettema, T. J. More than the sum of its parts: uncovering emerging effects of microbial interactions in complex communities. FEMS Microbiol. Ecol. fiae029 (2024).

68. Alseth, E. O., Custodio, R., Sundius, S. A. et al. The impact of phage and phage resistance on microbial community dynamics. PLoS Biol. 22, 1–25 (2024).

69. Castledine, M. & Buckling, A. Critically evaluating the relative importance of phage in shaping microbial community composition. Trends Microbiol. (2024).

70. Kruschke, J. K. Rejecting or accepting parameter values in bayesian estimation. Adv. Methods Pract. Psychol. Sci. 1, 270–280 (2018).

71. Haario, H., Laine, M., Mira, A. et al. DRAM: Efficient adaptive MCMC. Stat. Comput. 16, 339–354 (2006).

72. Gelman, A. & Rubin, D. B. Inference from iterative simulation using multiple sequences. Stat. Sci. 7, 457–472 (1992).

73. Vats, D. & Knudson, C. Revisiting the Gelman–Rubin diagnostic. Stat. Sci. 36, 518–529 (2021).

74. Geweke, J. Evaluating the accuracy of sampling-based approaches to the calculation of posterior moments. Staff Report 148, Federal Reserve Bank of Minneapolis (1991).

