## Supplementary Information for "Density-dependent feedback and higher-order interactions enable coexistence in phage-bacteria community dynamics"

for


#### S1 Mathematical modeling and inference

This section describes each of the classes of mechanistic ecological models used in this work, and details related to forward simulation and model-data fitting via Bayesian inference. First, we introduce baseline models to describe the interactions between a single host population and its virus followed by a scaled-up version of this model to describe dynamics across a community of bacteria and viruses. Finally, we introduce an additional emergent interaction for the community based on the accumulation of cellular debris. Following the model descriptions we provide technical details regarding Bayesian MCMC inference and model-data fitting.

##### S1.1 Pairwise SEIV model: phage–host pairwise dynamics

We model phage–bacteria ecological dynamics using a system of nonlinear coupled differential equations, where state variables track the density of susceptible bacteria ($S$), infected bacteria ($E$ and $I$) and virulent phage $V$. Susceptible bacteria are infected by phage with an effective adsorption rate $\phi$ and infections lead to lysis. We represent the latent period distribution between adsorption and lysis via a linear chain trick such that a host passes through $N_E$ different stages of infection ("exposed", $E^{(1)}$ to $E^{(N_E)}$) before reaching the final state ("infected", $I$) and lysing. We scale the transfer across infection stages such that it takes a newly infected host an average time of $\tau$, the latent period, to be lysed following an Erlang distribution whose variance is determined by $N_E$[17]. Once lysed, new virions are released with a burst size $\beta$. Table S2 provides details of the parameters. Throughout, we assume free phage virions may adsorb to any host cell, the system is not nutrient-limited, and that virus decay


[††]Mailing address: 105 Biological Sciences Building, 484 W 12th Ave, Columbus, OH 43210, USA

[‡‡]Mailing address: 3279 Biology-Psychology Building, 4094 Campus Dr, College Park, MD 20742, USA

is negligible on experimental time scales. We model the phage–host pairwise interaction as:

$$
\begin{aligned}
\dot{S} &= \overbrace{rS}^{\text{growth}} - \overbrace{\phi SV}^{\text{new infection}} \\
\dot{E}^{(1)} &= \overbrace{\phi SV}^{\text{new infection}} - \overbrace{\frac{N_E+1}{\tau}E^{(1)}}^{\text{delay}} \\
&\vdots \\
\dot{E}^{(N_E)} &= \overbrace{\frac{N_E+1}{\tau}E^{(N_E-1)}}^{\text{delay}} - \overbrace{\frac{N_E+1}{\tau}E^{(N_E)}}^{\text{delay}} \\
\dot{I} &= \overbrace{\frac{N_E+1}{\tau}E^{(N_E)}}^{\text{delay}} - \overbrace{\frac{N_E+1}{\tau}I}^{\text{lysis}} \\
\dot{V} &= \overbrace{\beta\frac{N_E+1}{\tau}I}^{\text{lysis}} - \overbrace{V\phi\left(S+\sum_i^{N_E}E^{(i)}+I\right)}^{\text{adsorption}}.
\end{aligned}
\tag{S1}
$$

#### S1.2 Pairwise SEIVD model: including infection attenuation

We include the potential for density-dependent attenuation of infection through the addition of a new state variable $D(t)$, which can be interpreted as a proxy for cellular debris. We model the probability of the virus infecting the host with a *Hill function*, given by $\frac{1}{1+\left(\frac{D}{D_c}\right)^2}$, where $D_c$ indicates the critical debris concentration at which the probability of viral adsorption to the bacterial strain $i$ falls to $\frac{1}{2}$. In the absence of debris ($D \ll D_c$) there is no inhibition to phage adsorption and in the limit of $D \gg D_c$ there are no new adsorption events to the $i$-th bacteria. The pairwise SEIVD model is given by,

$$
\begin{aligned}
\dot{S} &= \overbrace{rS}^{\text{growth}} - \overbrace{\frac{1}{1+\left(\frac{D}{D_c}\right)^2}}^{\text{debris-based attenuation}} \overbrace{\phi SV}^{\text{new infection}} \\
\dot{E}^{(1)} &= \overbrace{\frac{1}{1+\left(\frac{D}{D_c}\right)^2}}^{\text{debris-based attenuation}} \overbrace{\phi SV}^{\text{new infection}} - \overbrace{\frac{N_E+1}{\tau}E^{(1)}}^{\text{delay}} \\
&\vdots \\
\dot{E}^{(N_E)} &= \overbrace{\frac{N_E+1}{\tau}E^{(N_E-1)}}^{\text{delay}} - \overbrace{\frac{N_E+1}{\tau}E^{(N_E)}}^{\text{delay}} \\
\dot{I} &= \overbrace{\frac{N_E+1}{\tau}E^{(N_E)}}^{\text{delay}} - \overbrace{\frac{N_E+1}{\tau}I}^{\text{lysis}} \\
\dot{V} &= \overbrace{\beta\frac{N_E+1}{\tau}I}^{\text{lysis}} - \overbrace{\frac{1}{1+\left(\frac{D}{D_c}\right)^2}}^{\text{debris-based attenuation}} \overbrace{\phi V\left(S+\sum_i^{N_E}E^{(i)}+I\right)}^{\text{adsorption}}. \\
\dot{D} &= \overbrace{\frac{N_E+1}{\tau}I}^{\text{debris accumulation}}
\end{aligned}
\tag{S2}
$$

##### S1.3 Community SEIV model: Scaled-up community dynamics

In order to model population dynamics in communities with multiple interacting hosts and viruses, we scale up the system in Model A to consider the interacting terms for each individual strain of bacteria and phage. We use the index $i$ to refer to bacteria strains, $j$ for phage strains, and $ij$ for bacteria-phage pairs (e.g. the bacteria strain $i$ infected by phage strain $j$ is denoted $I_{ij}$). Each bacteria strain, phage strain, and bacteria-phage pair also has unique life-history traits (see Table S2 and Table S3 for strain names). We denote a binary interaction matrix $M$ ($M_{ij} = 1$ if host $i$ is infected by phage $j$ and zero otherwise). For the 9 different interactions present in the community we have 9 such values of burst sizes ($\beta_{ij}$), latent periods ($\tau_{ij}$), and adsorption rates ($\phi_{ij}$). The model dynamics are:

$$\begin{aligned}
\dot{S}_i &= \overbrace{r_i S_i}^{\text{growth}} - \overbrace{S_i \sum_j^{N_V} M_{ij}\phi_{ij}V_j}^{\text{new infection}} \\
\dot{E}_{ij}^{(1)} &= \overbrace{M_{ij}\phi_{ij}S_iV_j}^{\text{new infection}} - \overbrace{\frac{N_E^{ij}+1}{\tau_{ij}}E_{ij}^{(1)}}^{\text{delay}} \\
&\vdots \\
\dot{E}_{ij}^{(N_E^{ij})} &= \overbrace{\frac{N_E^{ij}+1}{\tau_{ij}}E_{ij}^{(N_E^{ij}-1)}}^{\text{delay}} - \overbrace{\frac{N_E^{ij}+1}{\tau_{ij}}E_{ij}^{(N_E^{ij})}}^{\text{delay}} \\
\dot{I}_{ij} &= \overbrace{\frac{N_E^{ij}+1}{\tau_{ij}}E_{ij}^{(N_E^{ij})}}^{\text{delay}} - \overbrace{\frac{N_E^{ij}+1}{\tau_{ij}}I_{ij}}^{\text{lysis}} \\
\dot{V}_j &= \overbrace{\sum_i^{N_H}\beta_{ij}\frac{N_E^{ij}+1}{\tau_{ij}}I_{ij}}^{\text{lysis}} - \overbrace{V_j\sum_i^{N_H} M_{ij}\phi_{ij}B_i}^{\text{adsorption}},
\end{aligned} \tag{S3}$$

where $B_i$ is the total density of bacteria strain $i$ across the susceptible, exposed, and infected classes:

$$B_i = S_i + \sum_j^{N_V}\sum_k^{N_E^{ij}} E_{ij}^{(k)} + \sum_j^{N_V} I_{ij}. \tag{S4}$$

#### S1.4 Community SEIVD model: Scaled-up community dynamics with infection attenuation

Similar to Model S1.2, we introduce a debris mediated infection attenuation model, controlled by the critical coefficient $D_{c_i}$ for each host $i$. The expanded model dynamics are:

$$
\begin{aligned}
\dot{S}_i &= \overbrace{r_i S_i}^{\text{growth}} - \overbrace{\frac{1}{1+\left(\frac{D}{D_{c_i}}\right)^2}}^{\text{debris-based attenuation}} \overbrace{S_i \sum_j^{N_V} M_{ij}\phi_{ij} V_j}^{\text{new infection}} \\
\dot{E}_{ij}^{(1)} &= \overbrace{M_{ij}\phi_{ij}V_j S_i}^{\text{new infection}} \overbrace{\frac{1}{1+\left(\frac{D}{D_{c_i}}\right)^2}}^{\text{debris-based attenuation}} - \overbrace{\frac{N_E^{ij}+1}{\tau_{ij}} E_{ij}^{(1)}}^{\text{delay}} \\
&\vdots \\
\dot{E}_{ij}^{(N_E^{ij})} &= \overbrace{\frac{N_E^{ij}+1}{\tau_{ij}} E_{ij}^{(N_E^{ij}-1)}}^{\text{delay}} - \overbrace{\frac{N_E^{ij}+1}{\tau_{ij}} E_{ij}^{(N_E^{ij})}}^{\text{delay}} \\
\dot{I}_{ij} &= \overbrace{\frac{N_E^{ij}+1}{\tau_{ij}} E_{ij}^{(N_E^{ij})}}^{\text{delay}} - \overbrace{\frac{N_E^{ij}+1}{\tau_{ij}} I_{ij}}^{\text{lysis}} \\
\dot{V}_j &= \overbrace{\sum_i^{N_H} \beta_{ij} \frac{N_E^{ij}+1}{\tau_{ij}} I_{ij}}^{\text{lysis}} - V_j \sum_i^{N_H} M_{ij}\phi_{ij} \overbrace{\frac{1}{1+\left(\frac{D}{D_{c_i}}\right)^2}}^{\text{debris-based attenuation}} \left( S_i + \sum_k^{N_E^{ij}} \sum_j^{N_V} E_{ij}^{(k)} + \sum_j^{N_V} I_{ij} \right) \\
\dot{D} &= \overbrace{\sum_i^{N_H} \sum_j^{N_V} \frac{N_E^{ij}+1}{\tau_{ij}} I_{ij}}^{\text{debris accumulation}}.
\end{aligned}
\tag{S5}
$$

#### S1.5 Simulating dynamics

We use MATLAB's non-stiff ordinary differential equation solver `ode45()` to numerically integrate the system of coupled differential equations. We set relative and absolute error tolerance to $10^{-8}$ and a non-negative criterion for our simulations. We convert all volumes to ml and times to hrs (see Table S2 for parameter units). Following experimental protocol, we simulate the one-step growth curves for the first 15 minutes numerically with experimentally measured initial conditions using Equation S1. Then we scale all the state variable densities by $10^{-2}$ due to a 100-factor dilution and simulate again with the same equation (Equation S1) for the duration of the experiment. For the community experiment, we set initial values for susceptible bacteria $S_i$ and free phage $V_j$ from the experimentally recorded value for each replicate and simulated for the experimental duration of 15 hours and 45 minutes. We obtain the total density $B_i$ for each strain of bacteria $i$ by summing across susceptible, exposed, and infected states (Equation S4). For phage, we report $V_j$ given that measurements are of free phage.

#### S1.6 Estimating the life-history parameters with Bayesian statistics

We use Bayesian MCMC to fit our mechanistic ecological models of phage-bacteria dynamics against the experimental datasets. The MCMC analyses are run with the MATLAB package *mcmcstat*, using a Delayed Rejection Adaptive Metropolis (DRAM) algorithm[44,71].

##### *S1.6.1 Bayesian MCMC convergence diagnostics*

To test for chain convergence we use the *Gelman-Rubin (GR)*[72,73] and *Geweke* diagnostic tests[74]. The GR test compares the inter-chain with the intra-chain variance and we choose a value of the test statistic $\hat{R}_{GR} < 1.1$ for all the chains after the burn-in phase as an acceptance criterion. For $M$ parallel chains of length $L$ each the GR test statistic $\hat{R}_{GR}$ for each parameter is given by

$$\hat{R}_{GR} = \sqrt{\frac{\frac{L-1}{L}W + \frac{B}{L}}{W}}, \tag{S6}$$

where $B$ and $W$ are the between-chain and within-chain sample variances respectively.

We use the Geweke test to detect if the chain has stabilized. We compare the means of different parts of the chain after the burn-in period to discard the values if the parameter chains show the signature of drift from their stable mean. In this study, we use the initial 0.1% of the chain and the final 0.5% of the chain and compute their means $m_A$ and $m_B$ and $0^{th}$ power spectral densities, $\hat{S}_A(0)$ and $\hat{S}_B(0)$ respectively. So our case the test statistic for each parameter should be

$$\hat{R}_{geweke} = \frac{m_A - m_B}{\sqrt{\frac{\hat{S}_A(0)}{0.1N} + \frac{\hat{S}_B(0)}{0.5N}}}. \tag{S7}$$

Both tests are utilized to measure the precision and accuracy of our inference protocol.

##### *S1.6.2 Fitting the one-step phage growth curves*

The one-step growth curves for each phage–host combination are parameterized with 5 parameters, $r, \phi, \beta, \tau$, and $N_E$. We use the observed free virus density time series in our model-data fitting, with other state variables being latent. Our likelihood function is defined as a product of Log-normal functions of data and model deviation at each of the observed data points, where the variance of the log-likelihood, $\sigma_{LL}$, is constant for all the data points. This reduces the log-likelihood function to $LL = -\frac{n}{2}\log(2\pi\sigma_{LL}^2) - \frac{ss}{\sigma_{LL}^2}$, where $ss$ is the sum of square error between model and the data, and $n$ is the number of data-points. In each case, the prior of the variance $\sigma_{LL}^2$ is specified by an inverse gamma distribution, with the hyperparameters (mean and variance) set through the errors from conventional trait estimates. We use data from all the time points and replicates with equal weights in the likelihood function.

For each of the one-step growth curves, we fit data points combined from all replicates. We use strictly positive Gaussian priors with their means set from the conventionally estimated values (in Table S4 ) and their standard deviations as $\beta_{sd} = 150(0, 700)$, $\phi_{sd} = 10^{-7}(10^{-10}, 10^{-6})$ ml/hr, $\tau_{sd} = 5(0.25, 5)$ hr, $N_{E_{sd}} = 100(5, 400)$, where truncation in each variable is denoted by (lower-bound,upper-bound). The prior of $\phi$, the adsorption rate, is Log-normal, whereas for the rest of the life-history parameters, it is a normal distribution. The prior covariance between parameters is set to zero. For $CV(\tau)$, we sample $N_E$, the number of compartments in our sequential infection model. See Fig S9 for the trace plots, prior distribution used and posterior distribution inferred, and convergence statistics for each of the parameter chains, using Gelman-Rubin tests. See Fig S10 for the details of the posterior covariance. We find that the latent period and burst size for a pair are positively correlated, whereas the adsorption rate is uncorrelated or weakly negatively correlated to the latent period and burst size, for most of our one-step experiments. A total of 10000 samples are collected for each case where the first 7000 samples are set as the burn-in. In each case, the intra-chain autocorrelation falls quickly within $\pm 0.2$ indicating proper mixing.

##### *S1.6.3 Fitting the 15.75-hour community experiment using scaled-up SEIV model*

We use a coordinate descent algorithm[45], to find the point-wise parameter estimates of the life-history traits (9 different sets of burst sizes, 9 different adsorption rates, 9 latent periods, and 5 host growth rates). We initialize the coordinate descent with an initial parameter estimate found by minimizing error across a bounded parameter space via Latin hypercube sampling with 500 samples. We chose the parameter bounds as follows, 50 to 700 virions/cell for burst sizes, $10^{-9}$ to $10^{-7}$ ml/hr for adsorption rates, 0.5 to 10 hrs for mean latent periods, 0.1 to 0.7 cells/hr for bacterial growth rates. The error function for the $k$-th replicate is given as the sum-of-squares of differences between the log-transformed model and data points,

$$\text{error function}_k = \sum_t^{N_T}\sum_i^{N_H}\left(\ln\hat{B}_i(t) - \ln B_{i,t,k}\right)^2 + \sum_t^{N_T}\sum_i^{N_V}\left(\ln\hat{V}_j(t) - \ln V_{j,t,k}\right)^2, \tag{S8}$$

where $\hat{B}_i(t), \hat{V}_j(t)$ are the simulated total host and phage densities at sample time $t$. The data points $B_{i,t,k}, V_{j,t,k}$ are the qPCR measurements of host $i$ and free phage $j$ respectively at time $t$ for the $k$-th replicate. We keep the coefficient of variation of the latent period the same as the one-step pairwise interactions. For $\phi$38:1 on CBA 18, where the one-step is not known we set the coefficient of variation of the latent period to 0.1.

##### S1.6.4 Fitting the initial community experiment time-series using the scaled-up SEIV model

Similar to the previous subsection, we use a coordinate descent algorithm to find representative point estimates for life history parameters over the first 6.4 hours of the community experiment after which viral-induced lysis outweighs bacterial growth (Table S9b). Next, we fit this initial phase with the SEIV model through the MCMC-based Bayesian inference and infer the distribution of the life-history traits. Similar to the one-step growth curve example, our Log-normal likelihood function treats each point independently for all replicates and the $\sigma_{LL}$ is sampled from an inverse $\gamma$ prior. In the SEIV model, the dynamics of CBA and PSA are completely uncoupled allowing us to perform the Bayesian inference separately on them. We run each chain for 50000 steps, discarding the first 20000 samples as burn-in. As this is a high-dimensional problem with an adaptation step of 100 samples, we also set our chain thinning to every 10 points. We achieve an acceptance rate of 68% and finally use 3000 working samples from the inferred joint posterior distribution.

##### S1.6.5 Fitting the 15.75 hrs community experiment with the SEIVD model

We use a strictly positive Gaussian prior with fixed means and variances for each of the parameters – burst sizes, mean of latent periods, and the host growth rates. We assume no prior covariance between these parameters. For the adsorption rate and critical debris concentration, we set up Log-Normal prior distributions, (see Fig. S12). We keep the coefficient of variation of the latent period (CV = $\frac{1}{\sqrt{N_E+1}}$) fixed to 0.07, for all the interactions for simplicity, as we observed that gave us the best fit from gradient descent based estimation[45]. The Bayesian inference scheme is exactly the same as described in the previous section S1.6.4. We run each chain for 50000 steps, discarding the first 20000 samples as burn-in. As this is a high-dimensional problem with an adaptation step of 100 samples, we also set our chain thinning to every 10 points. We achieve an acceptance rate of 72% and finally use 3000 working samples from the inferred joint posterior distribution. Fig S12 reports the inferred posterior distribution and Fig S13 reports the trace plots of the chains. For the posterior distributions, we use Matlab's `kde` (kernel density estimate) function to visualize the histogram for each parameter. Also in Fig S14 see the Gelman-Rubin and Geweke convergence statistics which remain less than 1.1 and greater than 0.9 for each of the chains, thus making the inference scheme acceptable with at least 90% level of significance. After thinning the intra-chain autocorrelation falls below a critical 0.2 for all the chains, indicating the chains are well mixed.

#### S1.7 Comparing Bayesian estimates of traits

To compare posterior estimates of a parameter between models, we computed the distribution of differences coming from 2 chains, A and B, for the same parameter $\theta$ defined as $\Delta = \theta_A - \theta_B$ from posterior samples. We defined the posterior probability of practical equivalence as $p_{\text{eq}} = P(|\Delta| < \varepsilon)$, with $\varepsilon$ a pre-specified Region Of Practical Equivalence (ROPE). We defined ROPE as 10% of the pooled standard deviation of the chains A and B combined. We classified evidence for differences based on $p_{\text{eq}}$. We considered the differences as strong when $p_{\text{eq}} < 0.0001$, as moderate differences when $0.0001 \leq p_{\text{eq}} \leq 0.01$, minor differences when $0.01 < p_{\text{eq}} < 0.05$, and inconclusive otherwise[70].

### S2 Additional experimental methods

#### S2.1 Primers

The forward and reverse primers for qPCR are shown in Table S7, along with the corresponding primer annealing temperatures.

#### S2.2 Effect of community spent media on pairwise infections

To test the effect of lysate on pairwise infections, we added the spent media from the community to all 9 fresh pairwise experiments found in Fig 1a. We filtered samples from the community at 12 hours with a 0.2$\mu$m filter and treated them with UV for 1.5 hrs to inactivate phages. We confirmed inactivity via a spot test. We added the samples back into fresh pairwise infections. We added 25$\mu$l of filtered, UV-treated sample, host strain of interest at $2 \times 10^6$ cells/ml, and phage of interest at $2 \times 10^5$ PFU/ml to 200$\mu$l total volume in a 96-well plate. We also included two controls: one without the phage, and one without the spent media sample. We measured growth by $OD_{600}$ every hour for a total of 16 hrs. The uninfected host densities increased in all the cases. However, with the addition of the 12-hour community spent media, infection attenuation is observed across all infection pairs, and the host density remains above the detection threshold (Fig. S5). In another control, spent media from 0 hr of community was added to fresh infections, and the slowdown of infections was not observed (data not shown).

#### S2.3 Burst size experiments on PSA strains

We combine PSA-HP1 phage with all the other 5 hosts (both PSA and CBA strains), at a MOI of 0.5 corresponding to each host. Similarly, in another experiment, we combine PSA-HS6 phage with all the other 5 hosts at a MOI of 0.5 for each host. We record the one-step growth curves of these phages on all the hosts (which we call the five-hosts-one-phage subcommunity). Then we use Model S1.1 to perform Bayesian inference on the phage one-step growth on this combined host system. We use the average of the conventionally measured parameter distribution as the priors of this inference. This allows us to obtain effective burst sizes, effective latent periods, and effective adsorption rates of the individual stain on the combination of hosts. We evaluate putative burst size shifts using a one-sided t-test. Our null hypothesis is that there is no difference in the mean burst sizes in the presence of all the hosts compared to the individuals. For PSA-HP1, the alternate hypothesis is that the burst size on all the hosts combined is greater than that on individual hosts.

#### S2.4 Resistance tests

Extra cell pellets from both infected and no phage control at 11.7 hrs, of the original pilot experiment that had been flash frozen and stored at -80°C were thawed. These samples were spread on plates and 15 colonies with CBA morphology were isolated and purified. These were then challenged with phage and possible resistance to phage infection determined by following growth via $OD_{600}$ every hour for 19 hrs. Identity of host strains was confirmed by the host range for CBA (where each host strain is infected by a different set of phages). We were not able to test PSA in this way due to extremely poor recovery of viable cells from frozen samples.

**Pairwise Experiments**

| **Type of experiment** | **Quantity measured** | **Models used** |
|---|---|---|
| Phage one-step growth for 8 phage host pairs (see main Fig. 2) | Phage density via plaque assays for each case | Pairwise SEIV model fitted |
| Phage adsorption assay (see main Fig. 2) | Adsorption rate of particular phage to host | Pairwise adsorption model |
| Multiple life cycle pairwise phage-host experiments (see main Fig. 3, SI Fig. S3) | Phage and infected host densities at 0 hrs, 3 hrs, and 15.75 hrs with phage-free infected host (control) | Pairwise SEIV and SEIVD model with parameters from one-step growth experiments |
| Pairwise phage-host experiments with added community spent media (see SI Fig S8) | Host densities via OD600 – phage free host, phage infected host (controls) and phage infected host with community spent media | NA |

**Community Experiments**

| **Type of experiment** | **Quantity measured** | **Models used** |
|---|---|---|
| 5 host community (uninfected control), (see main Fig. 1, SI Fig. S1) | Strain-resolved host densities every 35 minutes via qPCR for 15.75 hours | NA |
| 5 host 5 phage community (See main Fig. 5, SI Fig. S4, SI Fig. S5) | Strain-resolved host and phage densities every 35 minutes via qPCR for 15.75 hours | 1. Community SEIV model with pairwise parameters.<br>2. Community SEIV model with inferred parameters.<br>3. Community SEIVD model with pairwise traits and samples of $D_c$.<br>4. Community SEIVD model with inferred parameters |

**Sub-community Experiments**

| **Type of experiment** | **Quantity measured** | **Models used** |
|---|---|---|
| 5 host 1 phage subcommunity (for PSA only) (See SI Fig. S8) | Phage growth over time (via plaque assays) | Pairwise SEIV model (effective traits considered) |

**Table S1.** List of models and experiments used in this study.

Glossary of variables and units

| hyperparameter | description | units |
|---|---|---|
| $N_H$ | number of bacterial strains | - |
| $N_V$ | number of phage strains | - |
| $N_E^{ij}$ | number of exposed classes of bacteria $i$ via phage $j$ | - |
| **state variable** | **description** | **units** |
| $S_i$ | susceptible bacteria population $i$ | cells/ml |
| $E_{ij}^{(k)}$ | bacteria population $i$ exposed to phage population $j$ in the $k$th stage of infection ($0 \leq k \leq N_E^{ij}$) | cells/ml |
| $I_{ij}$ | bacteria population $i$ infected by phage population $j$ | cells/ml |
| $B_i$ | sum of bacteria $i$'s population (Equation S4) | cells/ml |
| $V_j$ | phage population $j$ | virions/ml |
| **parameter** | **description** | **units** |
| $r_i$ | bacteria $i$ growth rate | 1/hr |
| $M_{ij}$ | interaction for host $i$ - phage $j$ (boolean) | - |
| $\phi_{ij}$ | adsorption rate for host $i$ - phage $j$ | ml/hr |
| $\beta_{ij}$ | burst size for host $i$ - phage $j$ | virions/cell |
| $\tau_{ij}$ | latent period for host $i$ - phage $j$ | hr |
| $D_{c_i}$ | critical debris concentration for bacterial strain $i$ | dead cells/ml |

**Table S2.** Parameters, hyperparameters, and state variables for the phage–bacteria community model (Equation S3 and S5). See Table S3 for which bacteria and phage strains are assigned to which indices.

| $i$ | bacteria strain | $j$ | phage strain |
|---|---|---|---|
| 1 | CBA 4 | 1 | CBA $\phi$18:2 |
| 2 | CBA 18 | 2 | CBA $\phi$18:3 |
| 3 | CBA 38 | 3 | CBA $\phi$38:1 |
| 4 | PSA H100 | 4 | PSA-HP1 |
| 5 | PSA 13-15 | 5 | PSA-HS6 |

**Table S3.** Index assignment for the 5 bacteria and 5 phage strains. By convention, $i$ refers to bacteria strains and $j$ refers to phage strains. Model parameters with a double index $_{ij}$ refer to the pair of host $i$ and phage $j$.

| SL No | Phage | Host | r (cells/hr) | | $\phi$ ($10^{-8}$ ml/hr) | | $\tau$ (hr) | | $\beta$ (virions/cell) | | CV of $\tau$ |
|---|---|---|---|---|---|---|---|---|---|---|---|
| | | | C | B | C | B | C | B | C | B | C |
| 1 | $\phi$18:2 | CBA 18 | 0.24 | 0.25 | 2.1 | 1.9 | 1.85 | 1.87 | 91.2 | 126.6 | 0.082 |
| 2 | $\phi$18:3 | CBA 4 | 0.19 | 0.19 | 18.3 | 5.6 | 1.00 | 1.90 | 0.9 | 1.9 | 0.302 |
| 3 | $\phi$18:3 | CBA 18 | 0.24 | 0.24 | 13 | 12 | 1.42 | 2.03 | 27.3 | 63.8 | 0.076 |
| 4 | $\phi$38:1 | CBA 18 | 0.19 | NA | 18 | NA | $0.5-11$ | NA | $1-900$ | NA | NA |
| 5 | $\phi$38:1 | CBA 38 | 0.22 | 0.22 | 9.9 | 3.5 | 1.00 | 1.80 | 10.5 | 36.6 | 0.113 |
| 6 | HP1 | PSA H100 | 0.28 | 0.28 | 18 | 12.3 | 0.75 | 1.46 | 54.9 | 75.1 | 0.085 |
| 7 | HP1 | PSA 13-15 | 0.25 | 0..25 | 19 | 8.8 | 0.67 | 1.37 | 54.2 | 87.2 | 0.096 |
| 8 | HS6 | PSA H100 | 0.28 | 0.28 | 7.6 | 5.7 | 1.33 | 2.20 | 205.9 | 435.6 | 0.080 |
| 9 | HS6 | PSA 13-15 | 0.25 | 0.26 | 10 | 4.4 | 1.17 | 2.02 | 276.7 | 324.1 | 0.073 |

**Table S4.** Life-history parameters estimated from pairwise interacting phage–hosts single strain experiments from conventional analysis (C) and one-step growth curves of the viruses by Bayesian inference (B) – median values reported here; visualized in Fig. 2 with respective standard deviations. For $\phi$38:1 on CBA 18, where one-step growth experiments are not viable, a broad range of values is assumed from literature[46], where, $\tau$ can range from 0.5 to 11 hrs and burst size can range from 1 to 900 to simulate Fig. 5a.

| Phage | Host | $\beta$ (virions/cells) | $\phi$ (ml/hr) | $\tau$ (hr) | $r$ (cells/hr) | $D_c$ (cells/ml) |
|---|---|---|---|---|---|---|
| $\phi$18:2 | CBA 18 | 194.9 | $1.5 \times 10^{-8}$ | 1.7 | 0.22 | $6.19 \times 10^{6}$ |
| $\phi$18:3 | CBA 4 | 2.9 | $5.9 \times 10^{-8}$ | 2.9 | 0.18 | $5.06 \times 10^{6}$ |
| $\phi$18:3 | CBA 18 | 204.6 | $7.8 \times 10^{-8}$ | 2.7 | 0.22 | $6.19 \times 10^{6}$ |
| $\phi$38:1 | CBA 18 | 100.1 | $2.4 \times 10^{-8}$ | 2.3 | 0.22 | $6.19 \times 10^{6}$ |
| $\phi$38:1 | CBA 38 | 19.9 | $7.9 \times 10^{-8}$ | 1.9 | 0.29 | $11.26 \times 10^{6}$ |
| PSA-HP1 | PSA H100 | 525.4 | $6.1 \times 10^{-8}$ | 1.8 | 0.67 | $2.16 \times 10^{6}$ |
| PSA-HP1 | PSA 13-15 | 488.1 | $6.0 \times 10^{-8}$ | 2.3 | 0.53 | $1.61 \times 10^{6}$ |
| PSA-HS6 | PSA H100 | 60.6 | $1.1 \times 10^{-8}$ | 4.7 | 0.67 | $2.16 \times 10^{6}$ |
| PSA-HS6 | PSA 13-15 | 51.3 | $2.2 \times 10^{-8}$ | 1.9 | 0.53 | $1.61 \times 10^{6}$ |

**Table S5.** Median values of best fitted SEIVD model to community time series data. $r$ and $D_c$ are specified for each host, therefore is same across co-infecting phages.

| (a) Burst Size (virions/cells) | | | | | |
|---|---|---|---|---|---|
| Phage | Host | Pairwise SEIV model | Community SEIVD model | $p_{eq}$ | Decision |
| $\phi$18:2 | CBA 18 | 126.6 | 194.9 | 0.027 | Minor difference |
| $\phi$18:3 | CBA 4 | 1.9 | 2.9 | 0.001 | Moderate difference |
| $\phi$18:3 | CBA 18 | 63.8 | 204.6 | 0.004 | Moderate difference |
| $\phi$38:1 | CBA 38 | 36.6 | 19.1 | 0.070 | Inconclusive |
| PSA HP1 | PSA H100 | 75.1 | 525.5 | $< 10^{-4}$ | Strong difference |
| PSA HP1 | PSA 13-15 | 87.2 | 488.1 | $< 10^{-4}$ | Strong difference |
| PSA HS6 | PSA H100 | 435.6 | 60.6 | $< 10^{-4}$ | Strong difference |
| PSA HS6 | PSA 13-15 | 324.1 | 51.3 | $< 10^{-4}$ | Strong difference |
| (b) Latent period (hrs) | | | | | |
| Phage | Host | Pairwise SEIV model | Community SEIVD model | $p_{eq}$ | Decision |
| $\phi$18:2 | CBA 18 | 1.87 | 1.7 | 0.071 | Inconclusive |
| $\phi$18:3 | CBA 4 | 1.90 | 2.9 | 0.010 | Moderate difference |
| $\phi$18:3 | CBA 18 | 2.03 | 2.7 | 0.002 | Moderate difference |
| $\phi$38:1 | CBA 38 | 1.80 | 1.9 | 0.032 | Minor difference |
| PSA HP1 | PSA H100 | 1.46 | 1.8 | 0.025 | Minor difference |
| PSA HP1 | PSA 13-15 | 1.37 | 2.3 | $< 10^{-4}$ | Strong difference |
| PSA HS6 | PSA H100 | 2.20 | 4.7 | $< 10^{-4}$ | Strong difference |
| PSA HS6 | PSA 13-15 | 2.02 | 1.9 | 0.066 | Inconclusive |
| (c) Adsorption rate (in $10^{-8}$ ml/hr) | | | | | |
| Phage | Host | Pairwise SEIV model | Community SEIVD model | $p_{eq}$ | Decision |
| $\phi$18:2 | CBA 18 | 1.9 | 1.5 | 0.083 | Inconclusive |
| $\phi$18:3 | CBA 4 | 5.6 | 5.9 | 0.094 | Inconclusive |
| $\phi$18:3 | CBA 18 | 12 | 7.8 | 0.015 | Minor difference |
| $\phi$38:1 | CBA 38 | 3.5 | 7.9 | 0.048 | Minor difference |
| PSA HP1 | PSA H100 | 12.3 | 6.1 | 0.004 | Moderate difference |
| PSA HP1 | PSA 13-15 | 8.8 | 6.0 | 0.049 | Minor difference |
| PSA HS6 | PSA H100 | 5.7 | 1.1 | 0.045 | Minor difference |
| PSA HS6 | PSA 13-15 | 4.4 | 2.2 | 0.023 | Minor difference |

**Table S6.** Phage parameter (traits) comparison between pairwise SEIV model and community SEIVD model. We compared the posteriors of each parameter by setting a Region of Practical Equivalence (ROPE) and equivalent to 10% of the pooled standard of the combined chains. When $p_{eq} < 0.05$ the traits are different with a 95% confidence. Median values are noted for each case. See methods in section S1.7 and distributions in Fig. 6.

| Target | Name | F sequence | R sequence | $T_m$ (°C) |
|---|---|---|---|---|
| PSA H100 | H100 2 | GGTGAACTAATTCAATTGGGCGAT | ATCGGTGAGTGTCGAGGGT | 54 |
| PSA 13-15 | 13-15 1 | GAGTTTGTGTCGTTGGATCGT | CCCAACTAGTAAACCACCAATCA | 54 |
| CBA4 | 4 set A | TGTCGAGTTTTCTTTCAGTAGCGTG | ACGCCGCACTAATCATAGCCT | 62 |
| CBA18 | 18 set A | TTTTACGAGAACGCCATCTTTCCAC | TGATGTAAGAGGGTTGAGGGCT | 62 |
| CBA38 | 38 set B | CTAGCTCGTAACCCGTCAACCT | ATGGTGCTATTCACTTACTTCCTGC | 62 |
| PSA-HP1 | HP1 set 2 | TGAGCGATATGAGTGTCCGC | ACAAGCTTCCGACCAGAGTG | 62 |
| PSA-HS6 | HS6 set A | AACCCCGCCTTAGCAACTGT | TGATCAGCGGAGCCACTACG | 62 |
| $\phi$18:2 | 18:2 set 1 | AAAGGAATGCCCGGAGTCAG | GGGCCGCTGCATGATTAAAG | 63 |
| $\phi$18:3 | 18:3 set D | GGATAGGCCACGACGGAGAC | CCAATGCTTTCCGCATCCTTGA | 64 |
| $\phi$38:1 | 38:1 set B | AGACAGCCGACGATAACGCA | GCGTCGAAAGGTGTGAACCC | 62 |

**Table S7.** Primer sequences and annealing temperatures ($T_m$) used for qPCR for each bacterial strain and phage.

| Phage | Host | From where | $\beta$ (virions/cell) | $\beta$ on all hosts | $p_{eq}$ | Decision |
|---|---|---|---|---|---|---|
| PSA-HP1 | PSA H100 | pairwise SEIV inference | 75.1 | 136.7 | 0.0302 | Minor differences |
| PSA-HP1 | PSA H100 | community SEIVD inference | 525.5 | 136.7 | $< 10^{-4}$ | Strong differences |
| PSA-HP1 | PSA 13-15 | pairwise SEIV inference | 87.2 | 136.7 | 0.0432 | Minor differences |
| PSA-HP1 | PSA 13-15 | community SEIVD inference | 488.1 | 136.7 | $< 10^{-4}$ | Strong differences |
| PSA-HS6 | PSA H100 | pairwise SEIV inference | 435.6 | 247.0 | 0.0182 | Minor differences |
| PSA-HS6 | PSA H100 | community SEIVD inference | 60.6 | 247.0 | 0.0002 | Moderate differences |
| PSA-HS6 | PSA 13-15 | pairwise SEIV inference | 324.1 | 247.0 | 0.0416 | Minor differences |
| PSA-HS6 | PSA 13-15 | community SEIVD inference | 51.3 | 247.0 | 0.0002 | Moderate differences |

**Table S8.** Summary of ROPE test for comparing burst sizes of the phages on all the hosts (in one phage five hosts subcommunity) vs a single host. Median burst sized noted for reference.

| Phage | Host | $\beta$ (virions/cells) | $\phi$ (ml/hr) | $\tau$ (hr) | $r$ (cells/hr) |
|---|---|---|---|---|---|
| (a) whole 15.75 hours of the experiment | | | | | |
| $\phi$18:2 | CBA 18 | 478.8 | $1.26 \times 10^{-9}$ | 1.5 | 0.072 |
| $\phi$18:3 | CBA 4 | 203.7 | $6.29 \times 10^{-9}$ | 2.5 | 0.077 |
| $\phi$18:3 | CBA 18 | 124.1 | $1.24 \times 10^{-8}$ | 3.2 | 0.072 |
| $\phi$38:1 | CBA 18 | 183.3 | $2.99 \times 10^{-9}$ | 4.3 | 0.072 |
| $\phi$38:1 | CBA 38 | 191.8 | $4.13 \times 10^{-8}$ | 3.9 | 0.059 |
| PSA-HP1 | PSA H100 | 227.3 | $8.09 \times 10^{-8}$ | 1.0 | 0.472 |
| PSA-HP1 | PSA 13-15 | 888.2 | $1.51 \times 10^{-8}$ | 6.3 | 0.465 |
| PSA-HS6 | PSA H100 | 421.9 | $3.25 \times 10^{-8}$ | 10.6 | 0.472 |
| PSA-HS6 | PSA 13-15 | 761.9 | $4.40 \times 10^{-9}$ | 2.6 | 0.465 |
| (b) first 6.4 hours of the experiment | | | | | |
| $\phi$18:2 | CBA 18 | 416.4 | $1.99 \times 10^{-9}$ | 1.8 | 0.129 |
| $\phi$18:3 | CBA 4 | 174.4 | $1.57 \times 10^{-7}$ | 3.2 | 0.125 |
| $\phi$18:3 | CBA 18 | 164.3 | $2.02 \times 10^{-8}$ | 3.4 | 0.129 |
| $\phi$38:1 | CBA 18 | 123.1 | $4.66 \times 10^{-9}$ | 2.2 | 0.129 |
| $\phi$38:1 | CBA 38 | 112.2 | $2.56 \times 10^{-8}$ | 4.0 | 0.213 |
| PSA-HP1 | PSA H100 | 317.4 | $3.39 \times 10^{-8}$ | 3.3 | 0.527 |
| PSA-HP1 | PSA 13-15 | 753.3 | $8.99 \times 10^{-9}$ | 5.1 | 0.572 |
| PSA-HS6 | PSA H100 | 474.2 | $2.30 \times 10^{-8}$ | 3.3 | 0.527 |
| PSA-HS6 | PSA 13-15 | 389.6 | $1.07 \times 10^{-9}$ | 5.8 | 0.572 |

**Table S9.** Community life-history parameters inferred through the SEIV model fitted to the experiment using a coordinate gradient descent algorithm. SEIV community model is not an accurate representation of the community dynamics, so we caution against treating those estimates as realistic. This table is only provided for the sake of completion.

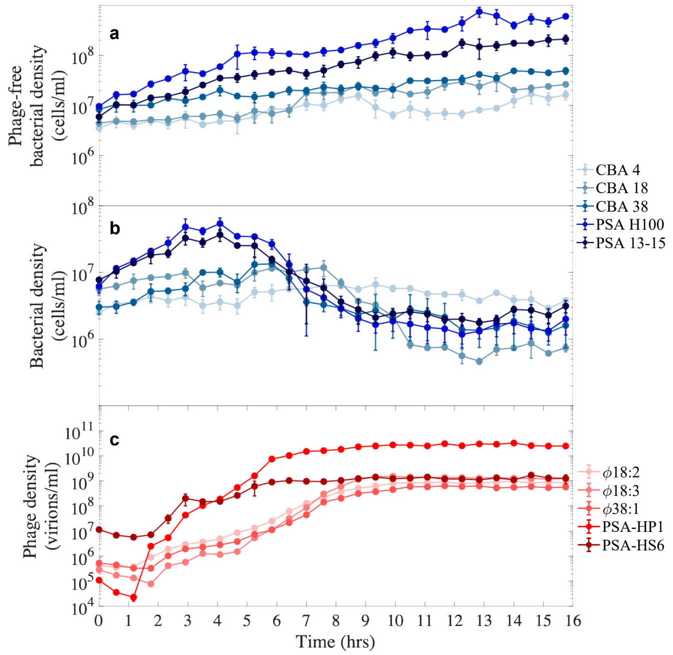


**Figure S1. Mean and standard deviations of phage-host population dynamics in the community and control. (a)** Strain-wise dynamics for the phage-free five host community. **(b-c)** Host and phage dynamics, respectively, for the five host five phage community.

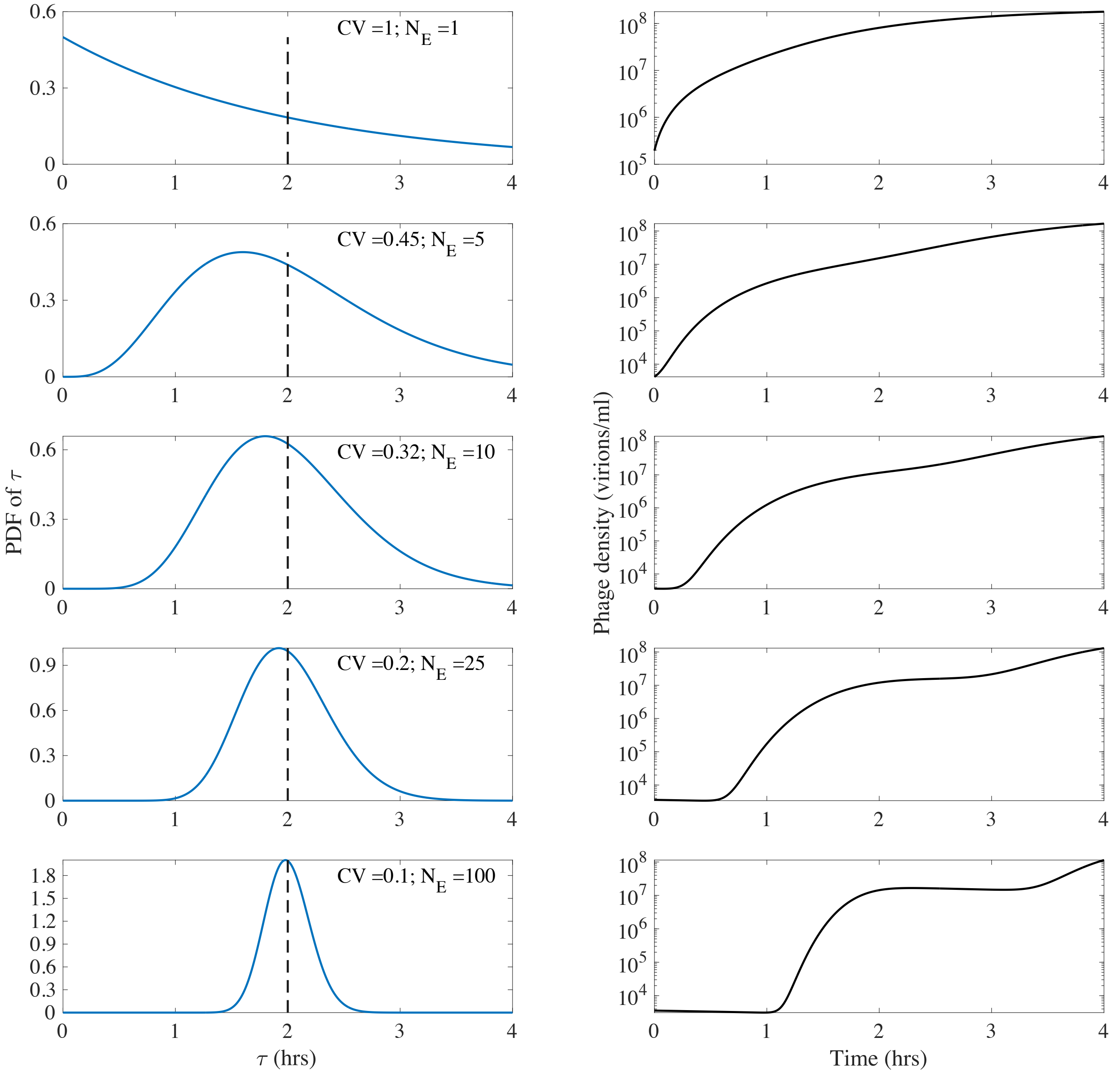


**Figure S2.** **Latent period distributions for different variations around mean.** Latent period of a host-phage modeled by a gamma distribution. Gamma distributions with different coefficient of variations tuned by the number of compartments is shown in the left, and the corresponding simulated viral trajectories shown in the right (simulation parameters: initial host density = $10^8$ host cells/ml, initial MOI = 0.1, $\beta = 200$ virions/cell, $\tau = 2$ hours, $\phi = 1.3 \times 10^{-8}$ ml/hr, $r = 0.2$ cells/hr). For number of exposed compartments, $N_E = 1$, the latent period corresponds to an exponential distribution, whereas with higher $N_E$ the distribution becomes sharper around the mean. For $N_E \to \infty$ the distribution converges to a Dirac delta function.

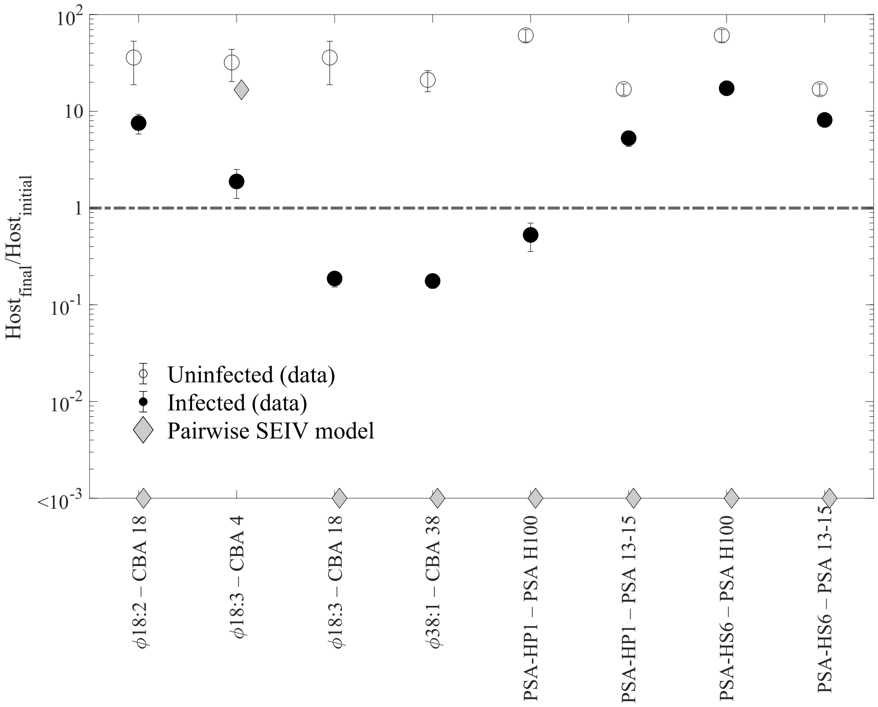


**Figure S3. Ratios of final to initial host densities.** Each experiment is performed for 15.75 hrs, representing multiple life cycles. Ratios of qPCR measured final (at 15.75 hrs) to initial (at 0 hrs) host densities for the 8 pairs of infection (infected in solid circles, control in white circles) and pairwise SEIV model parameterized by pairwise parameters from Fig. 2 (gray diamonds). Constant abundance is represented by the dashed line.

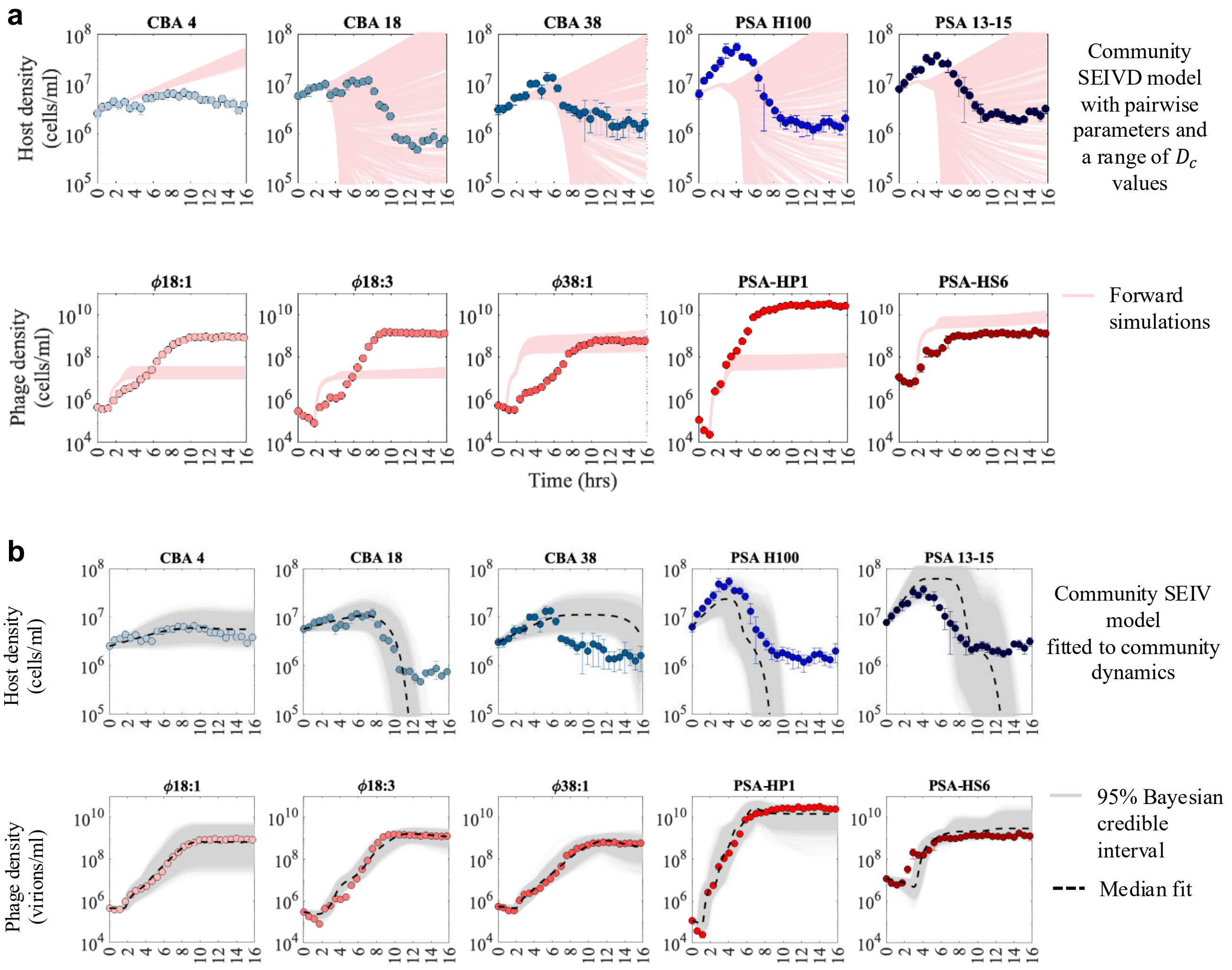


**Figure S4.** **Modeling only infection attenuation or change in ecological traits, but not both. (a)** The community SEIVD model parameterized by pairwise parameters of Fig. 2 and critical debris coefficients $D_{c_i}$ sampled from Log-Uniform distributions with a range $10^5 - 10^7$. (b) The scaled-up SEIV model fitted to the first 6.4 hours of the experiment successfully captures the initial part of the dynamics and predicts the later phage densities accurately and predicting selective bacterial strain elimination. The SEIV model is unable to fit bacterial dynamics for the whole duration (see Supplementary Information Fig S11). The fits from 95% credible intervals of the inferred parameter posteriors are plotted in gray, and median fits are shown with dashed lines.

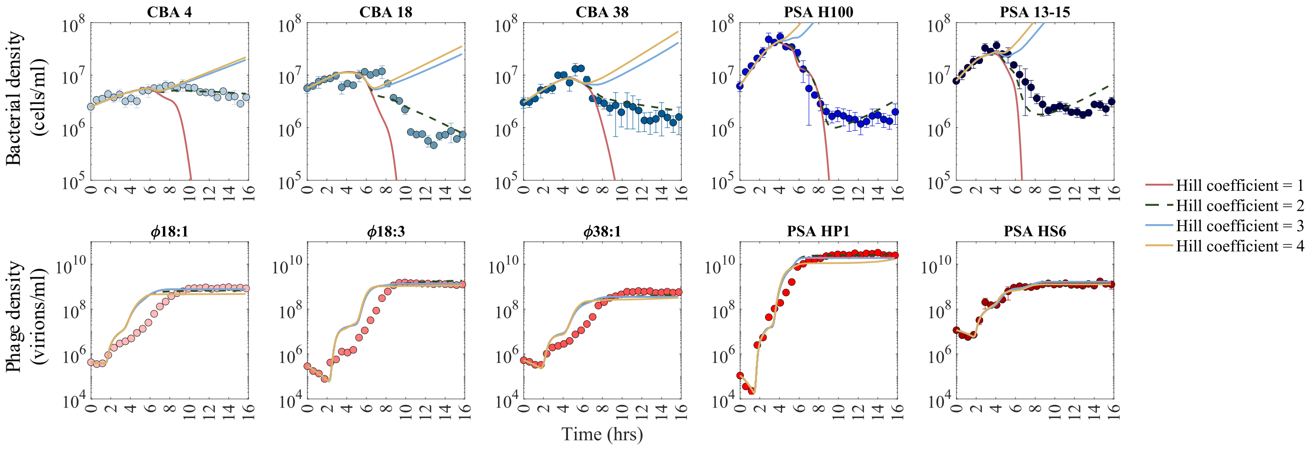


**Figure S5. Variation of Hill coefficient.** Community SEIVD model, parameterized by inferred median best fit values, with different Hill coefficients. The dashed line shows a Hill coefficient of 2 which has been used throughout the paper.

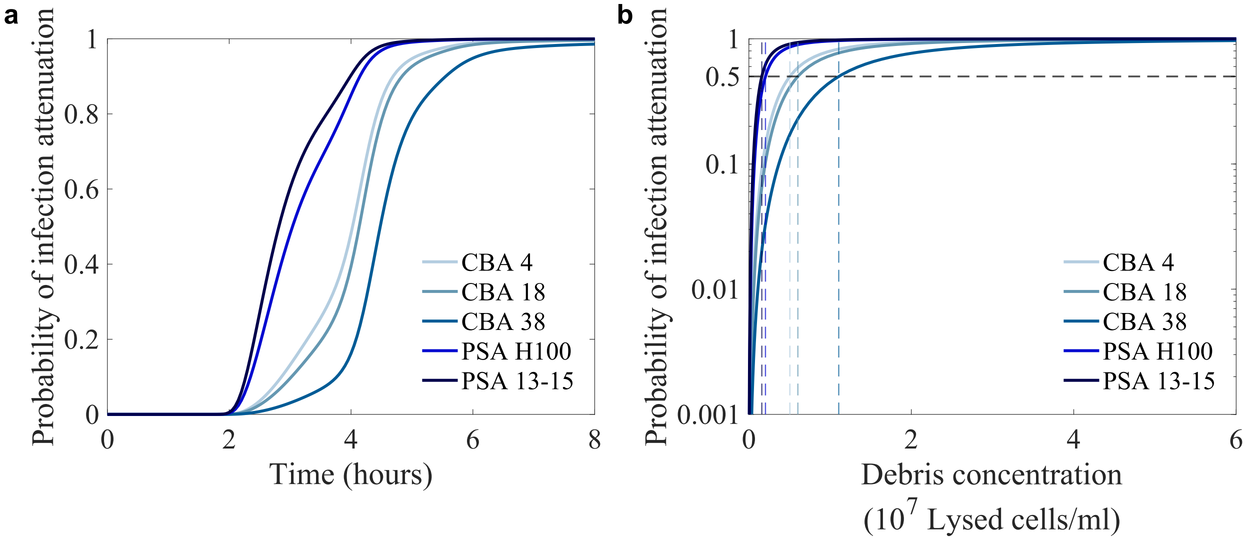


**Figure S6. Hill function driven attenuation of infection.** The time series trajectory for accumulated debris inferred from our model S1.4 (a) The probability of infection attenuation, which we define as $1 - \frac{1}{1+(D/D_{c_i})^2}$, shown for the PSA and CBA species, where the effect is much stronger in PSA. (b) The probability of infection attenuation increases with increasing concentration of debris in the system reaching a critical value of 0.5 for critical concentration $D_{c_i}$ of each strain (dashed vertical lines). The inferred median $D_{c_i}$ values are tabulated in Table S5.

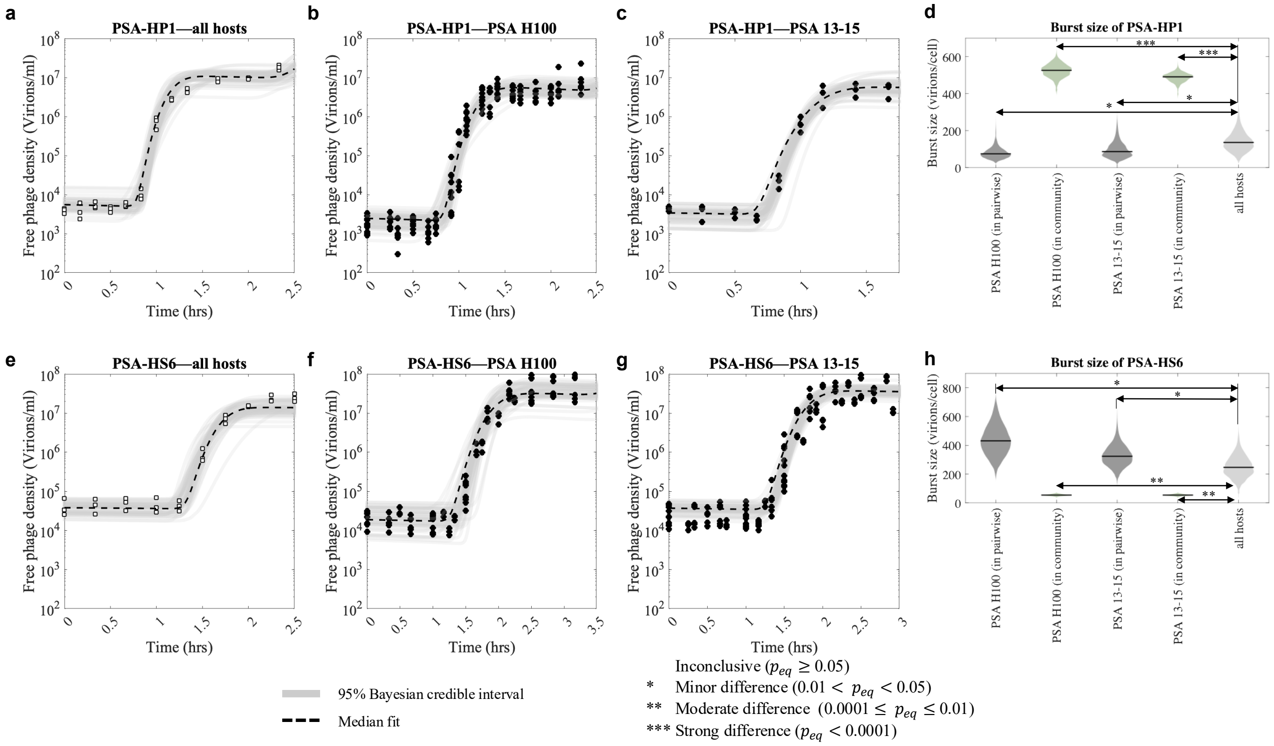


**Figure S7. Context-dependent PSA burst size.** One-step growth curves for PSA phages on single hosts and five hosts combined. (a-c) PSA-HP1 and (e-g) PSA-HS6. Here (b-c) and (f-g) are the same data and fits from Fig 2b. (d) and (h) show the burst sizes of HP1 and HS6 respectively inferred from community (green circle), 5 hosts combined subcommunity (open square) and pairwise experiments (black circle). Bayesian test of equivalence[70] (methods in Supplementary Information section S1.7) reveals that the effective burst size of HP1 with all hosts is less than the community values, whereas HS6 on all hosts is less than corresponding community values. Minor statistical difference is observed between pairwise values with individual hosts vs. all hosts combined; moderate differences were found between all host combined burst size and community values. See summary is Table S8.

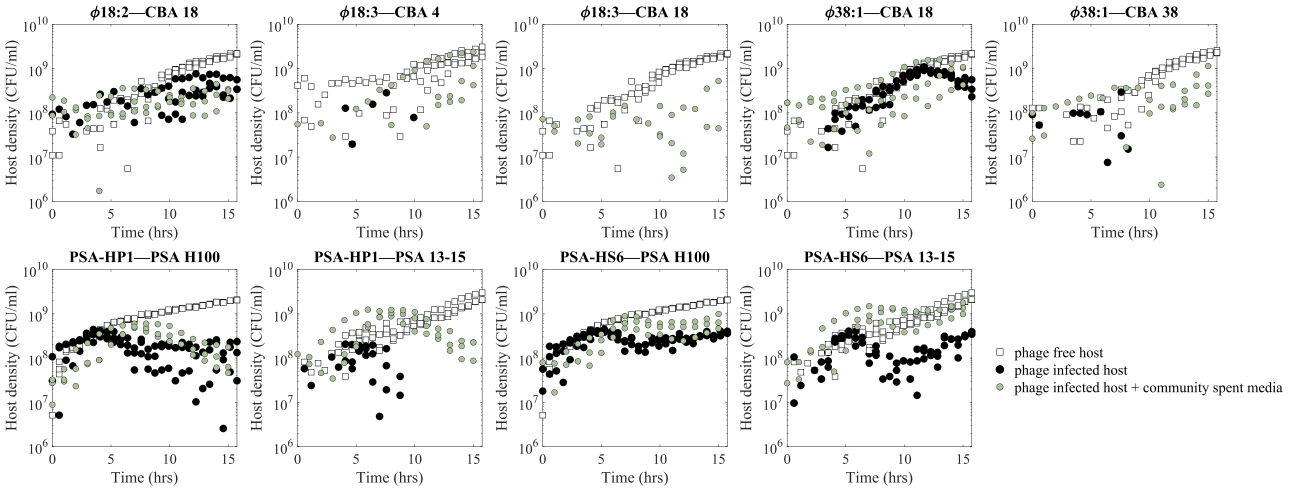


**Figure S8. Spent media added to fresh pairwise infections.** The spent media from the 12 hrs time point of the community is filtered and UV-treated and added to fresh pairwise infections (green circles). Their $OD_{600}$ is measured and plotted after calibrating it to CFU/ml along with phage-free control (squared) and phage-infected control (black circles). The cases where the data is not shown mean the OD600 values went below the level of detection.

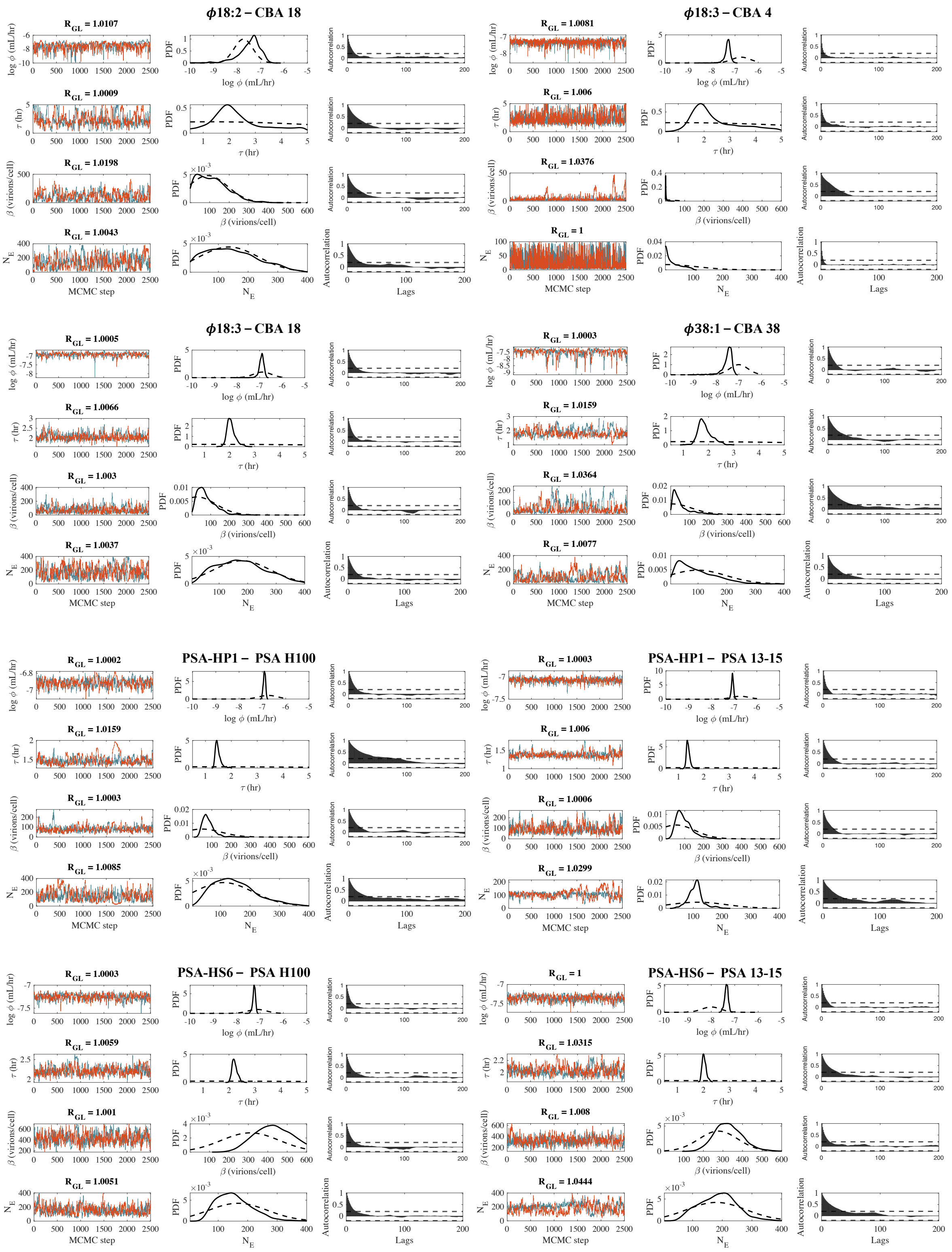


**Figure S9. One-step growth curve statistics and convergence:** The trace plots for two parallel MCMC runs, along with values of Gelman-Rubin $\hat{R}_{GR}$ statistics are shown along with the prior (dashed lines) and inferred posterior distributions (solid lines) for the trait parameters of each of the 8 virus-host pairs labeled in Fig 1. Magnitude of the chain autocorrelation rapidly falls within an acceptable range between $\pm 0.2$ indicating proper mixing.

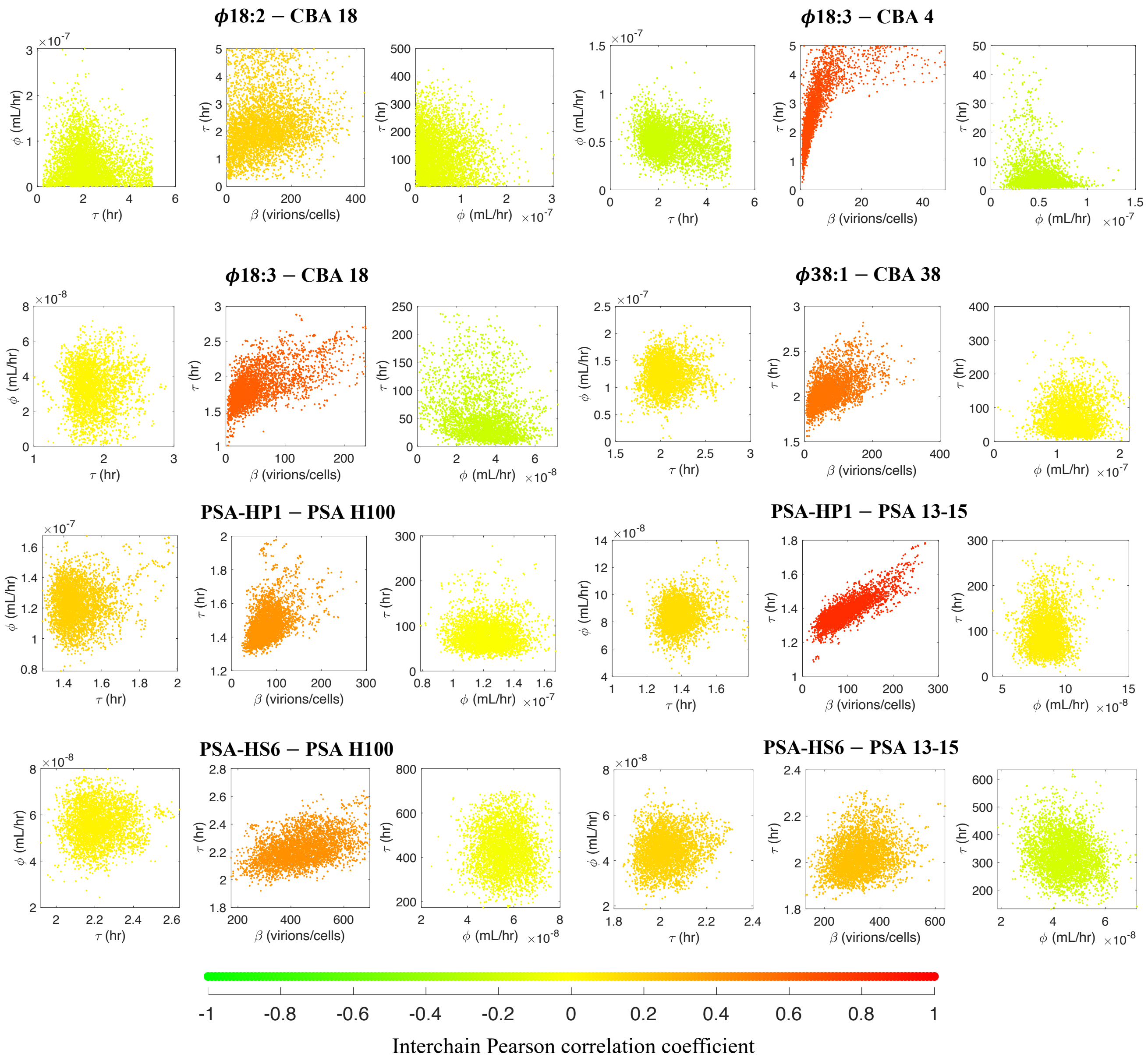


**Figure S10.** **Examples of parameter covariance in one-step models**: The covariance plots from the inferred joint posterior distributions for $\phi - \tau$, $\tau - \beta$, $\tau - \phi$ are shown for the 8 one-step growth curves, where each of the plots is color-coded according to their inter-chain Pearson correlation coefficient.

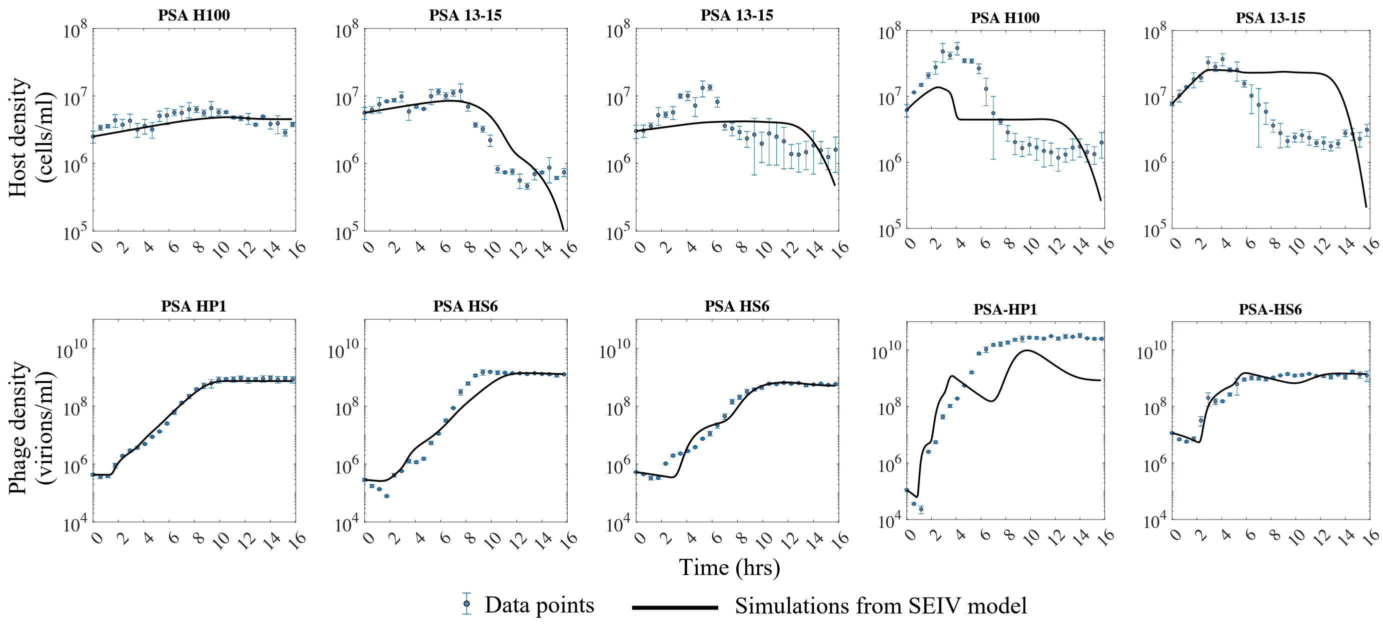


**Figure S11.** SEIV model fails to recapitulate the population dynamics when fitted across the whole duration of the experiment, with parameters in Tab. S9a.

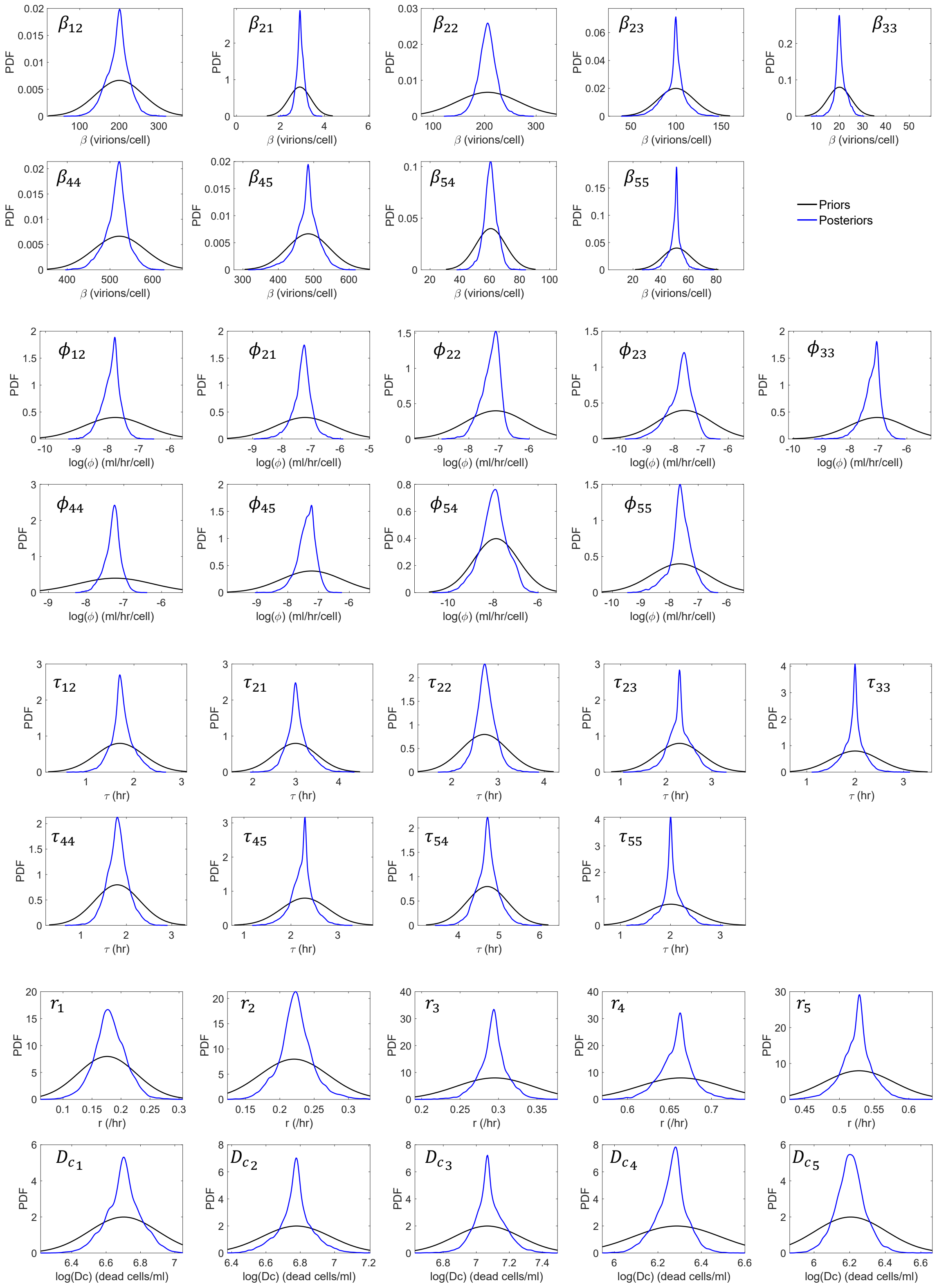


**Figure S12.** Priors and Posteriors distribution inferred from Bayesian modeling of the community time series data using the scaled up SEIVD model shown in black and blue lines respectively. $\phi$ and $D_c$ are shown in the log scale. See indices in Table S3.

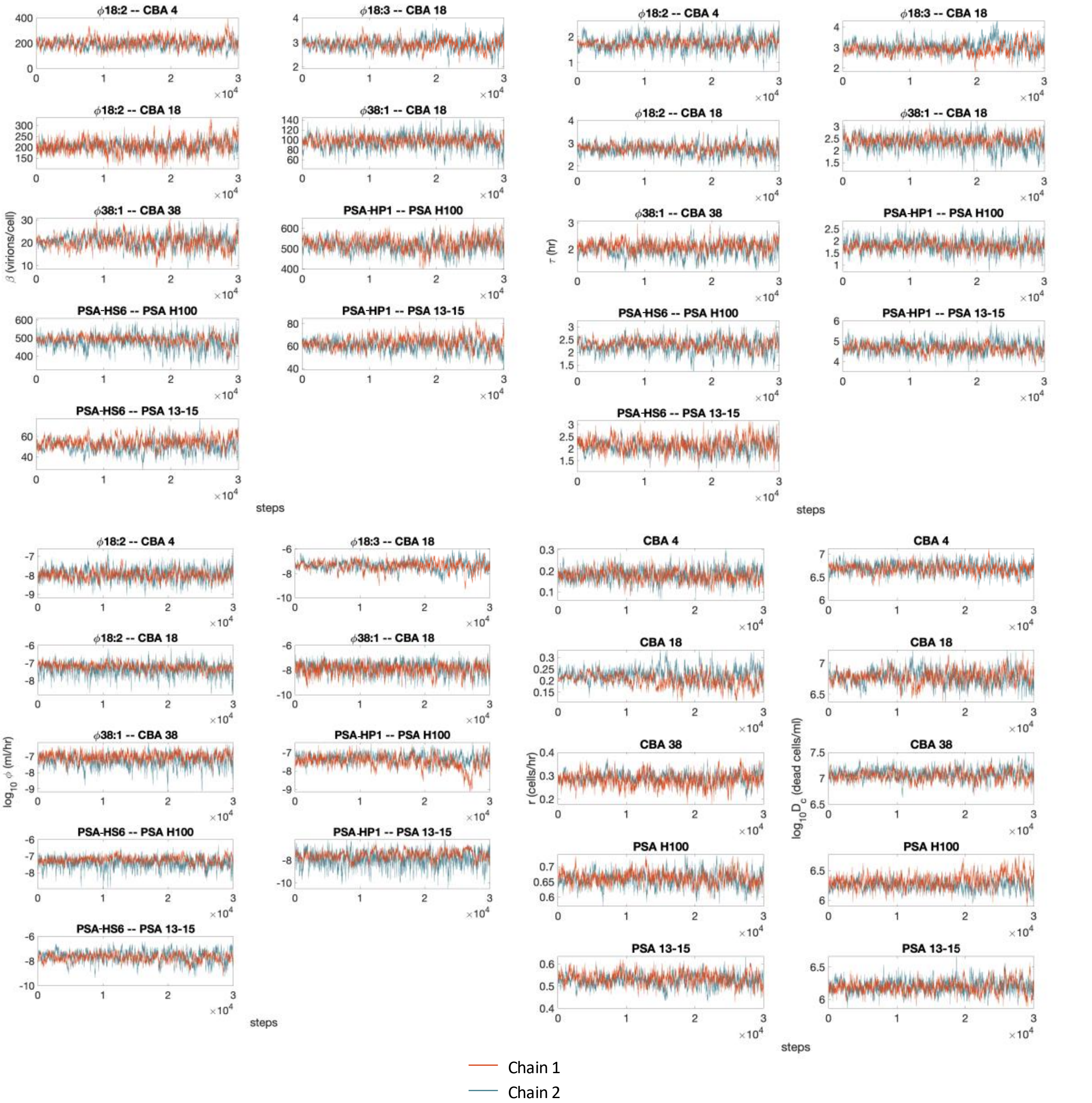


**Figure S13.** Traceplots of the life-history parameters shown for inference with SEIVD model (see Sec. S1.4) for two parallel runs in two different colors after removing the burn-in period of 20000 samples.

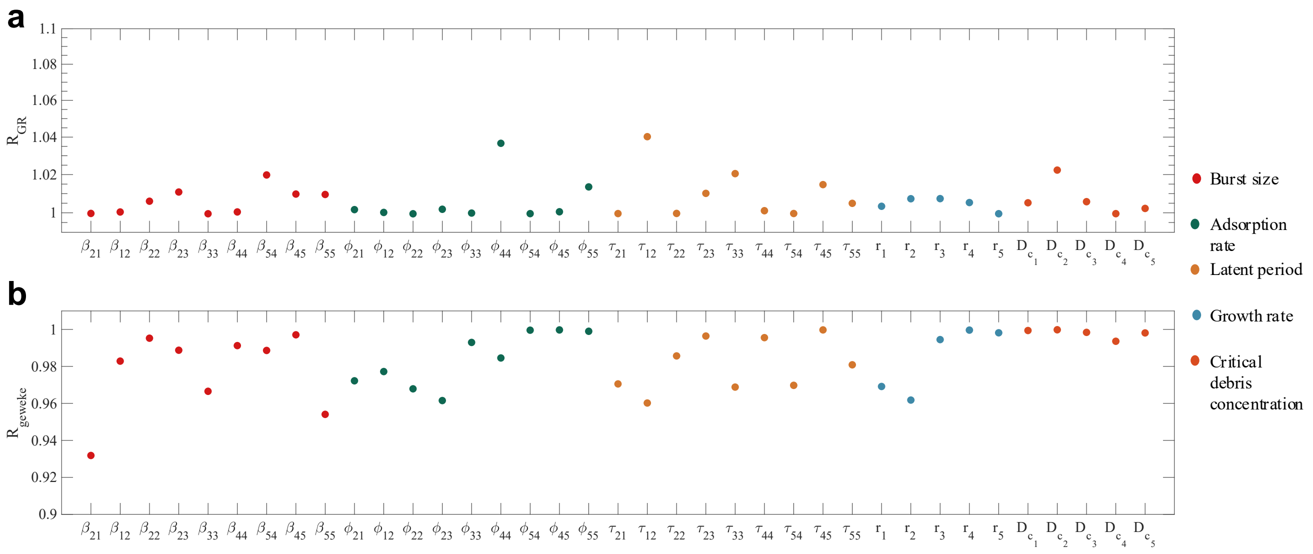


**Figure S14.** The results for the convergence tests shown for the two test statistics namely Gelman-Rubin (for 2 parallel chains) and Geweke in (a) and (b) respectively for the five growth rates ($r_i$), nine pairs of burst sizes ($\beta_{ij}$), adsorption rates ($\phi_{ij}$) and latent periods ($\tau_{ij}$) and the debris critical constant ($D_{c_i}$). The posterior distribution converges to a stationary state, as $R_{GR}$ is less than 1.1 and $R_G$ is greater than 0.9 for all the chains individually. See indices in Table S3.

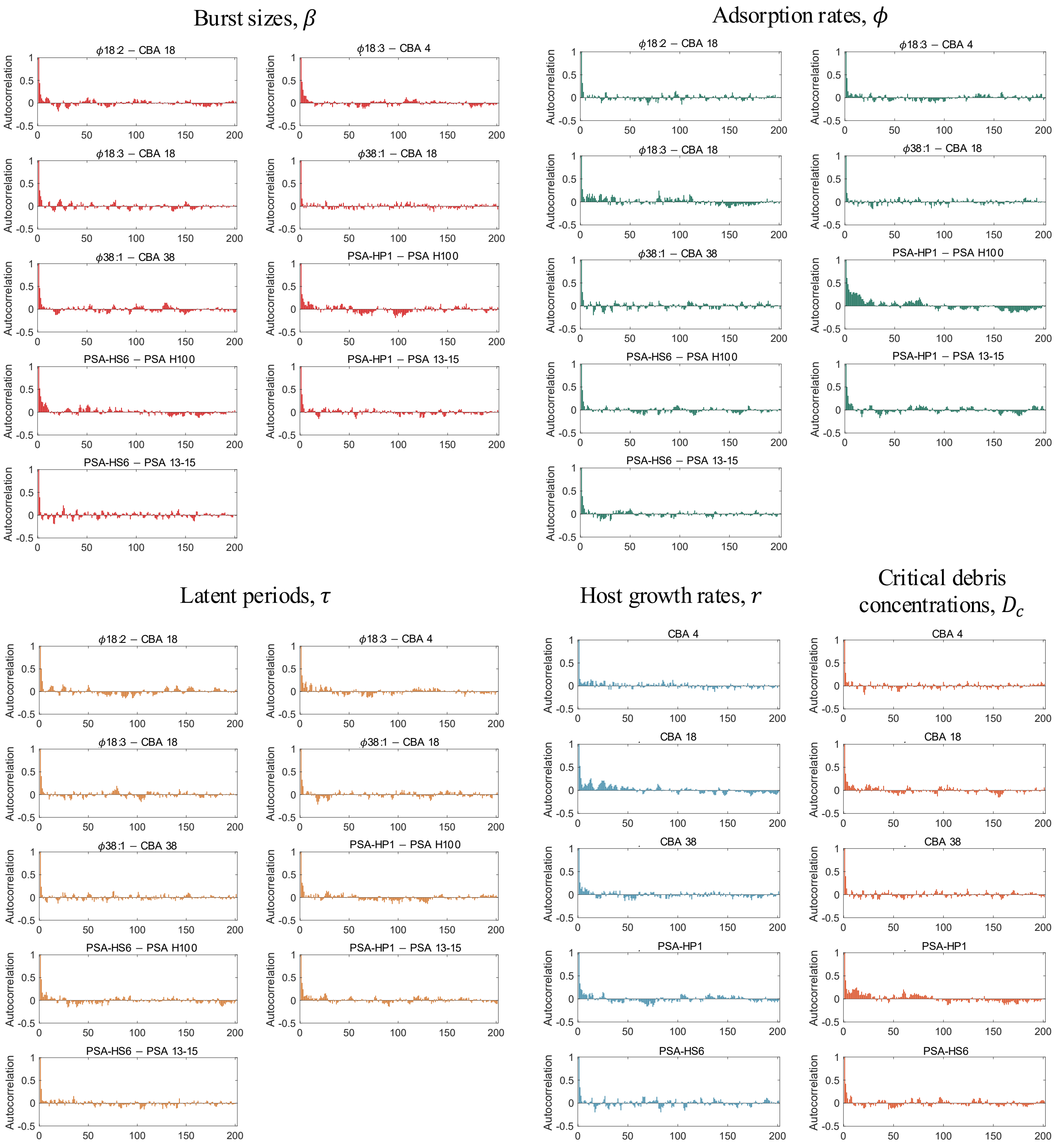


**Figure S15.** Intra-chain auto-correlations as obtained from the MCMC chains, fitting the community time series data with the scaled-up SEIVD model. We use a chain thinning of every 10 points and the absolute autocorrelation values rapidly fall below a 0.2 acceptable cutoff (dashed lines), indicating proper mixing.
